# Integrated transcriptomic correlation network analysis identifies COPD molecular determinants

**DOI:** 10.1101/764852

**Authors:** Paola Paci, Giulia Fiscon, Federica Conte, Valerio Licursi, Jarrett Morrow, Craig Hersh, Michael Cho, Peter Castaldi, Kimberly Glass, Edwin K. Silverman, Lorenzo Farina

## Abstract

Chronic obstructive pulmonary disease (COPD) is a heterogeneous and complex syndrome. Network-based analysis implemented by SWIM software can be exploited to identify key molecular switches - called “switch genes” - for disease. Genes contributing to common biological processes or define given cell types are frequently co-regulated and co-expressed, giving rise to expression network modules. Consistently, we found that the COPD correlation network built by SWIM consists of three well-characterized modules: one populated by switch genes, all up-regulated in COPD cases and related to the regulation of immune response, inflammatory response, and hypoxia (like *TIMP1*, *HIF1A*, *SYK*, *LY96*, *BLNK* and *PRDX4*); one populated by well-recognized immune signature genes, all up-regulated in COPD cases; one where the GWAS genes *AGER* and *CAVIN1* are the most representative module genes, both down-regulated in COPD cases. Interestingly, 70% of *AGER* negative interactors are switch genes including *PRDX4*, whose activation strongly correlates with the activation of known COPD GWAS interactors *SERPINE2*, *CD79A*, and *POUF2AF1*. These results suggest that SWIM analysis can identify key network modules related to complex diseases like COPD.

## Introduction

Chronic obstructive pulmonary disease (COPD) is a devastating lung disease characterized by progressive and incompletely reversible airflow obstruction. Like many other common diseases, COPD is a heterogeneous and complex syndrome influenced by both genetic and environmental determinants and is a major cause of morbidity and mortality throughout the world. Cigarette smoking is a major environmental risk factor for COPD, but the substantial heritability of COPD indicates an important role for genetic determinants as well [Zhou 2013]. Although multiple genetic loci for COPD have been identified by genome-wide association studies (GWAS), the key genes in those regions are largely undefined. Various contributors to COPD pathogenesis have been also suggested, including protease-antiprotease imbalance, oxidant-antioxidant imbalance, cellular senescence, autoimmunity, chronic inflammation, deficient lung growth and development, and ineffective lung repair. However, the pathobiological mechanisms for COPD remain incompletely understood [Silverman 2018].

COPD susceptibility, like other complex diseases, is rarely caused by a single gene mutation, but is likely influenced by multiple genetic determinants with interconnections between different molecular components. Studying the effects of these interconnections on disease susceptibility could lead to improved understanding of COPD pathogenesis and the identification of new therapeutic targets. Previous efforts to identify the network of interacting genes and proteins in COPD have included protein-protein interaction (PPI) network studies. McDonald and colleagues [McDonald 2014] used dmGWAS software to identify a consensus network module within the PPI network based on COPD GWAS evidence. Sharma and colleagues [Sharma 2018] started with “seed” genes based on well-established COPD GWAS genes or Mendelian syndromes that include COPD as part of the syndrome constellation with a random walk approach to build a COPD network module of 163 proteins.

Alternative approaches aiming to gain key insights into the genes driving the underlying disease molecular machinery are based on gene expression data. Typically, expression levels for each gene are compared between sample groups, and the genes that pass certain statistical cutoffs are selected as the “gold standard list" for further interpretation and validation. Recently, this kind of data analysis has been widely used to identify gene expression differences in lung tissue and blood between COPD cases and controls. Moving forward, in order to identify COPD causal genes, transcriptomic approaches can be complementary to genetic studies, as illustrated by studies relating differential gene expression to GWAS loci and to genetically predicted gene expression [Sakornsakolpat 2019, Morrow 2018, Morrow 2017]. However, in most cases, conclusions from differential expression studies are drawn using within-experiment data, thus claims of specificity are relative to the control groups used for reference, potentially leading to inaccurate interpretation [Crow 2019]. The reasons for this lack of specificity, especially in a highly heterogeneous and complex disease like COPD [Agusti 2014], can be multifarious. Among technical limitations, it is worth noting that economic restrictions typically limit the number of expression profiling experiments to a relatively small number of observations, thus preventing the identification of slight but significant changes [Häupl 2004]. Moreover, gene expression differences may be observed only in specific cell types and/or at specific stages of disease/process development. Among conceptual limitations, it is well-known that multiple cellular signaling pathways may impact the expression of the same gene making it difficult to identify the affected pathway from observing its expression changes [Häupl 2004]. In addition, cells may use many other mechanisms to regulate proteins besides changing the amount of mRNA, so these genes may remain constitutively expressed in the face of varying protein concentrations [Häupl 2004]. To overcome these limitations, it is necessary to complement differential expression analysis with other more sophisticated methodologies able to refine the “gold standard list” of differentially expressed genes (DEGs) gaining more specificity in the prediction of disease-associated genes.

Among others, popular approaches that start where DEG analysis ends are based on the construction of a co-expression network using, for example, Pearson correlation as a similarity index. Currently, two of the most promising algorithms for gene expression networks are SWIM (SWItchMiner) [Paci 2017] and WGCNA (Weighted Correlation Network Analysis) [Zhang 2005]. Both of them use the correlation structure to construct a gene-gene similarity network, divide the network into modules (groups of genes with similar expression), and identify “driver” genes in modules (WGCNA) or intra-modules (SWIM). Morrow and colleagues [Morrow 2017] used WGCNA to identify a network module differentially expressed in COPD that was related to B lymphocyte pathways. However, previous correlation-based network analyses in COPD have not used methods that can identify key molecular switches for disease [Mcdonough 2017, Morrow 2017, Chang 2016, Ezzie 2012].

As matter of fact, WGCNA considers only the right tail (i.e., positive correlation between gene pairs) of the correlation distribution. To date, the left tail (i.e., negative correlation between gene pairs) of the correlation distribution, and the interpretation of negative edges within a complex network representation of functional connectivity, has largely been ignored. Instead, it is well-known that the human genome is pervasively transcribed [ENCODE 2007], yet at any given spatial/temporal state a cell generally uses only a fraction of its gene functions. Thus, there are likely crucial roles of negative regulation to save cells from activation of specific pathways and cell functions in response to specific external stimuli or physiological and/or pathological changes. As an example, microRNAs are now universally recognized as key negative regulators in many intracellular processes as well as in cancer development and progression [Calin 2006, Zhou 2015].

The strength of the SWIM methodology is to emphasize the importance of negative regulation by explicitly considering the left tail of the correlation distribution. The main property of the driver genes identified by SWIM, called “switch genes”, is to be primarily anti-correlated with their partners in the correlation network: when switch genes are induced their interaction partners are repressed, and *vice versa*.

Here, we applied SWIM to lung tissue gene expression data from two well-characterized COPD case-control populations to study the differences between lung samples from normal subjects (represented by smokers with normal spirometry) and COPD cases. We used the dataset with a larger number of lung tissue samples (i.e., GSE47460 with 219 COPD cases and 108 controls) as the “training set” for running SWIM and the dataset of Morrow and colleagues [Morrow 2017] with a smaller number of samples (i.e., GSE76925 with 111 COPD cases and 40 controls) as the “test set” for validating the results.

We found that the COPD correlation network built by SWIM software consists of three modules, of which one includes multiple switch genes and is significantly enriched in pathways like: B cell receptor signaling pathway, NF-kappa B (NF-κB) signaling pathway, hypoxia, regulation of inflammatory response, regulation of immune response, collagen fibril organization, regulation of TGFB production, and extracellular matrix organization. We hypothesized that the SWIM approach would both support known pathways and provide evidence for novel pathways in COPD pathogenesis.

## Results

### COPD correlation network

The network-based analysis implemented by the SWIM software (see Materials and Methods section) was exploited to identify disease genes and network modules associated with COPD status by using the GSE47460 dataset (training set) containing microarray gene expression profiling of lung tissue samples from 219 COPD cases and 108 controls [Peng 2016, Anathy 2018].

Starting from 17530 genes, we obtained 2097 significantly differentially expressed genes (DEGs) at a 1% false discovery rate (FDR) [Benjamini-Hochberg 1995] (Figure 1 and Supplementary Table 1). We found 1358 DEGs (65%) down-regulated in COPD cases and the remaining 739 DEGs (35%) up-regulated (Figure 1a). Among DEGs, we found 145 genes located within genomic regions (+/− 1 Mb from the top SNP) previously identified as containing genome-wide significant associations to COPD [Sakornsakolpat 2019] (Supplementary Table 1).

**Figure 1.**
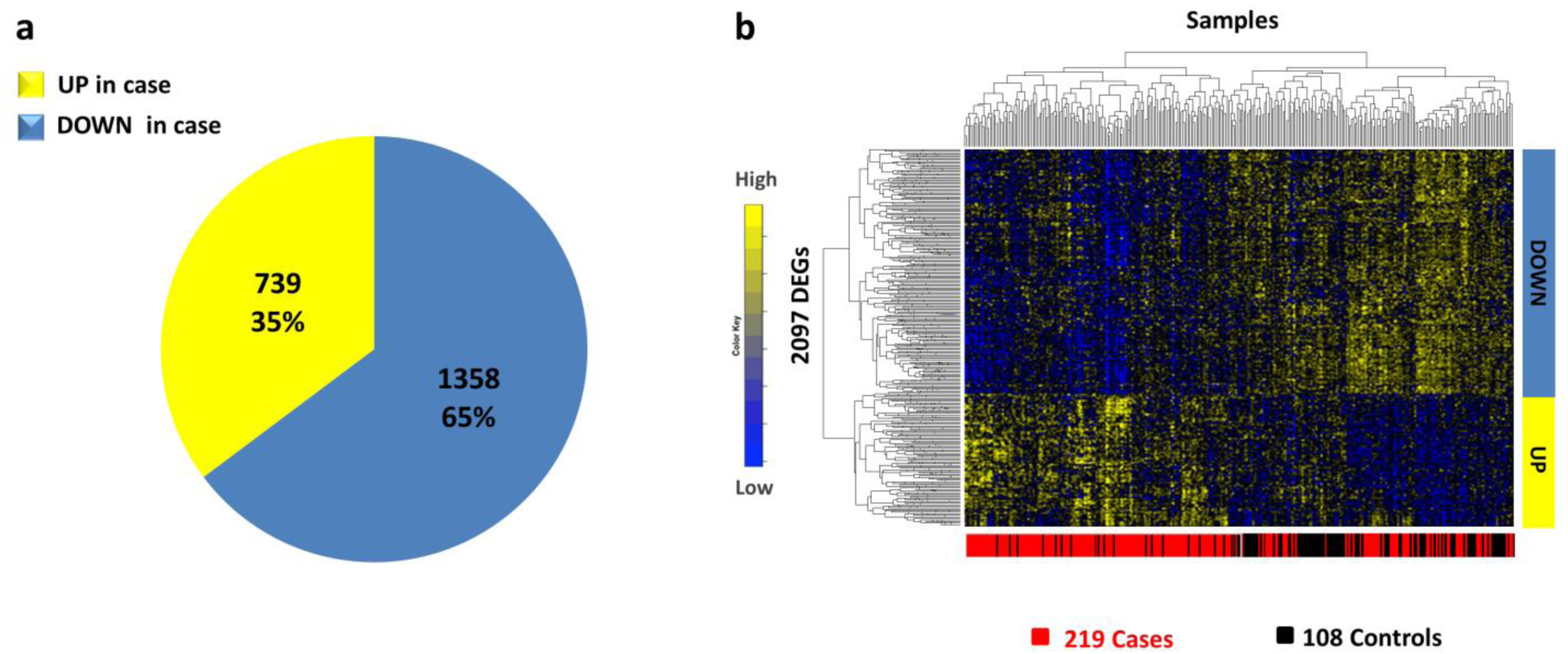
Differentially expressed genes in lung tissue samples. (a) Pie chart represents the percentages of DEGs that are up-/down-regulated in COPD cases in comparison to control subjects, based on 1% false discovery rate. (b) Heatmap represents DEGs clustered according to genes (rows) and samples (columns) by using one minus the Pearson correlation as distance. Colors represent different expression levels increasing from blue to yellow.

In order to check if the number of GWAS genes included in the DEGs is more than expected by chance, we randomly selected 2097 genes (i.e., the number of DEGs) from the original list of 17530 genes and repeated this procedure 1000 times. Then, the number of the 145 GWAS genes included in the DEGs was zscore-normalized and the p-value for the given z statistics was calculated; the p-value of 0.2 indicates that the number of differentially expressed GWAS genes is equal to what expected by chance. This observation is in accordance with the results obtained in [Morrow 2017], where the authors showed that COPD GWAS genes were not differentially expressed in lung tissue samples.

The DEG matrix of 2097 rows (DEGs) and 327 columns (219 COPD cases + 108 control samples) was used as input to SWIM in order to build the COPD gene correlation network based on the Pearson correlation coefficient, where a threshold is set for the absolute value of the minimum correlation coefficient necessary to draw an edge (see Materials and Methods section). For the COPD correlation network, we set the correlation threshold equal to 0.57, which corresponded to the 98^th^ percentile of the entire correlation distribution (Supplementary Figure 1).

The obtained COPD correlation network encompasses 1665 nodes and 52513 edges. The most highly connected hub is *EMP2* that codes a tetraspan protein of the PMP22/EMP family regulating cell membrane composition. It is down-regulated in COPD cases (Fold-change = 0.68, FDR = 1.1 ∙ 10^−7^) and is located in a genome-wide significant COPD GWAS region with a COPD-associated top SNP about 86 Kb from the transcription start site.

### Module identification in the COPD correlation network

In order to detect the community structure of the network, SWIM used the k-means clustering algorithm, which partitions *n* objects (here, network nodes) into a predefined number *N* of clusters (modules). The quality of clustering was evaluated by minimizing the Sum of the Squared Error (SSE), depending on the distance of each object to its closest centroid. As a distance measure, SWIM used:

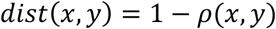

where ρ(*x*, *y*) is the Pearson correlation between expression profiles of nodes x and y. A reasonable choice of the number of clusters is suggested by the position of an elbow in the SSE plot (named “scree plot”) computed as a function of the number of clusters (see Materials and Methods section). The COPD correlation network consisted of 3 modules or clusters, varying in size from 190 genes in module 1, 1411 genes in module 2, and 64 genes in module 3 (Figure 2a).

In order to check the quality of the k-means clustering algorithm implemented by SWIM, we grouped genes with correlated expression profiles into modules by using complete linkage hierarchical clustering coupled with the correlation-based dissimilarity *dist*(*x*, *y*) = 1 − ρ(*x*, *y*). We compared detected modules with the ones obtained by SWIM with the k-means method, and we found that cluster 1 and cluster 3 are well separated meaning that their cluster detection is highly robust with respect to the clustering algorithm used (Supplementary Figure 2).

**Figure 2.**
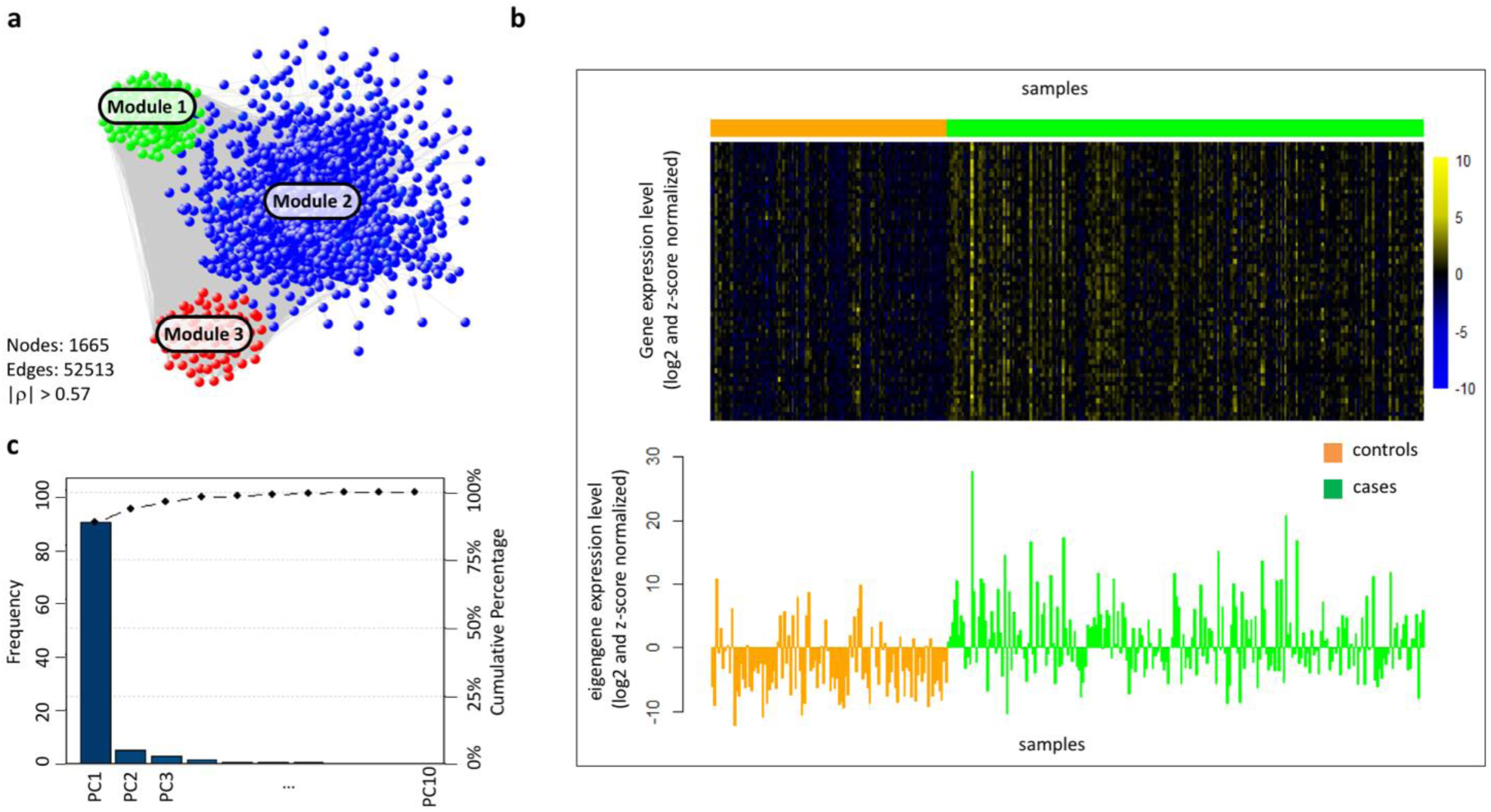
COPD correlation network and module eigengene. (a) COPD correlation network where nodes are DEGs and a link occurs between them if the absolute value of the Pearson correlation coefficient between their expression profiles exceeds the correlation threshold (|r| > 0.57). Groups of nodes sharing the same color represent gene modules obtained by k-means clustering. (b) [UPPER] Heatmap representing genes of module 3 (rows) across samples (columns). Colors represent different expression levels increasing from blue to yellow. Gene expression data are log2-transformed and z-score normalized. [BOTTOM] Bar plot of the expression levels of module 3 eigengene (y-axis) across samples (x-axis). Gene expression data are log2-transformed and z-score normalized. (c) The percent variability explained by each principal component (PC) computed for module 3, known as a Pareto chart, contains both bars and a line graph, where individual values are represented in descending order by bars, and the line represents the cumulative total value. The left y-axis represents the percentage of the data variance explained by each PC, the right y-axis represents the cumulative distribution, and the x-axis represents the PCs that are able to explain 100% of the cumulative distribution. PC1 represents the module eigengene and explains about 90% of the data variance.

### Summarizing the profiles of the COPD modules

To summarize the gene expression profiles of a given module in the COPD co-expression network, we used the module eigengene (Figure 2b) defined as the first principal component of a given module [Langfelder 2008]. We found that the first principal component across all modules is able to explain more than 85% of the data variance in each module, i.e. 96.7%, 95.4%, 88.8 %, in module 1, module 2, and module 3, respectively (Figure 2c). Thus, the module eigengene can be considered a representative gene able to condense each module into one profile. In light of this, we found that the eigengenes of module 1 and module 2 are both down-regulated in COPD cases (p-value = 5.8 ∙ 10^−14^ and p-value = 3.6 ∙ 10^−16^, respectively), whereas the eigengene of module 3 is up-regulated in COPD cases (p-value = 2.9 ∙ 10^−13^), providing a general idea of the overall expression trend of each module.

Then, for each gene in a given module, the module membership can be computed as the correlation between its expression profile and the module eigengene [Langfelder 2008]. We found high correlations within modules 1 and 3, as expected, with the mean module membership, or mean correlation of gene expression with the module eigengene, equal to 0.71 and 0.67, respectively. However, the mean module membership of module 2 is lower, confirming the result obtained with the hierarchical clustering algorithm (Supplementary Table 3).

The first two genes with the highest membership in module 1 are *CAVIN1* and *AGER*, both down-regulated in COPD cases (Fold-change = 0.8 and FDR = 1.8 ∙ 10^−5^; Fold-change = 0.65 and FDR = 1.8 ∙ 10^−5^, respectively). *CAVIN1* encodes a protein that enables the dissociation of paused ternary polymerase I transcription complexes from the 3’ end of pre-rRNA transcripts. This protein regulates rRNA transcription by promoting the dissociation of transcription complexes and the reinitiation of polymerase I on nascent rRNA transcripts. This protein also localizes to caveolae at the plasma membrane and is thought to play a critical role in the formation of caveolae and the stabilization of caveolins. *AGER*, one of the most down-regulated genes in COPD cases, encodes the advanced glycosylation end product (AGE) receptor, which is a member of the immunoglobulin superfamily of cell surface receptors. It is a multiligand receptor, and besides AGE, interacts with other molecules implicated in homeostasis, development, and inflammation (Figure 3). Interestingly, *AGER* is one of the most well-known candidate genes located in a significant COPD GWAS region with a non-synonymous SNP (located about 2 Kb from the transcription start site), which has been associated with multiple COPD-related phenotypes and COPD affection status [Kim 2015, Li 2018].

**Figure 3.**
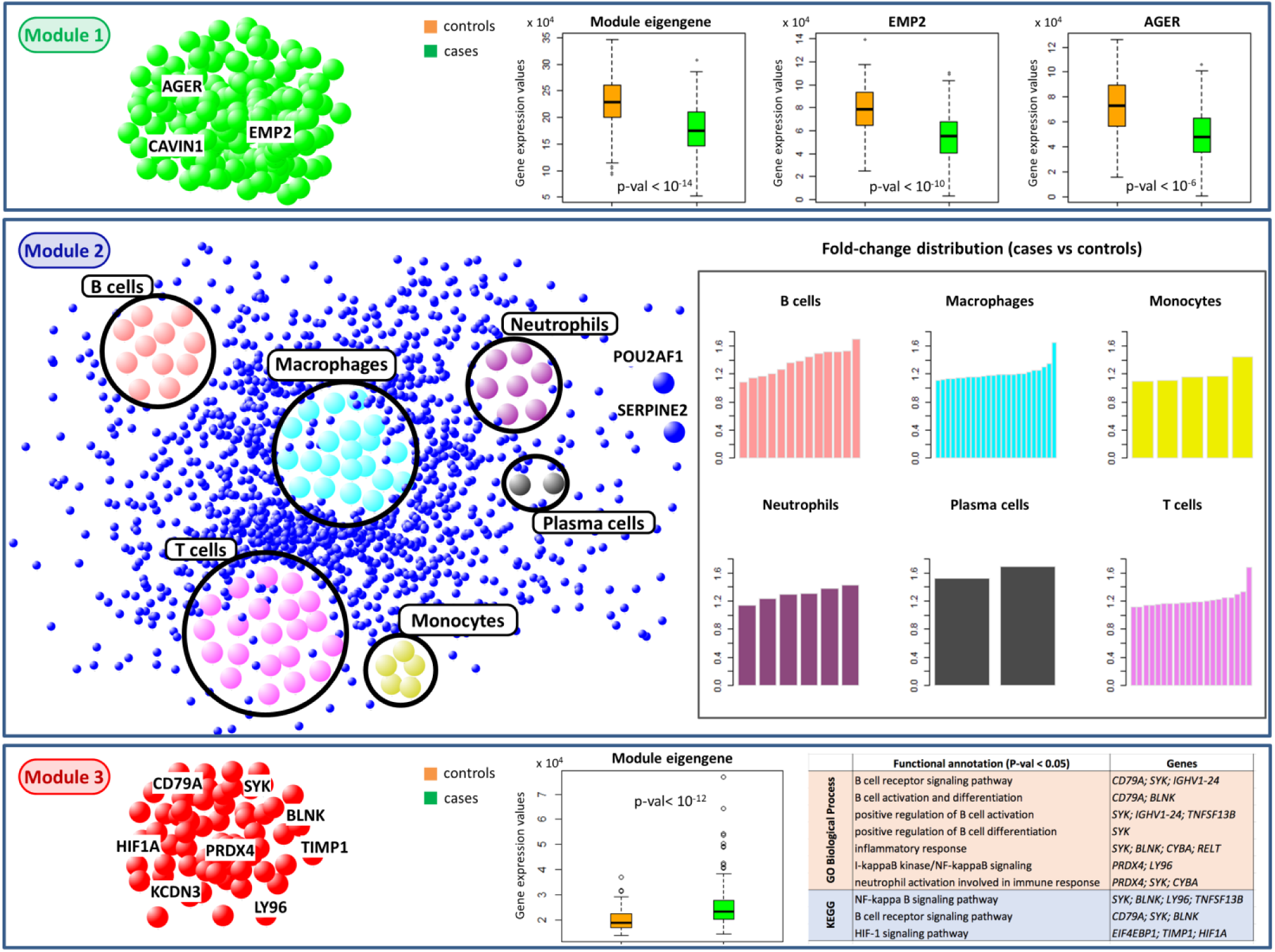
Module characterization in COPD network. The three boxes represent the three modules obtained by k-means clustering from the COPD correlation network. In each module, genes of interest or immune cell populations are highlighted. From top to bottom: boxplots in controls (orange boxes) and COPD cases (green boxes) of the module 1 eigengene and of the GWAS genes with the highest module 1 membership; bar plots, for each immune cell population included in module 2, of the fold-change values of the marker genes in that immune cell population; boxplot in controls (orange boxes) and COPD cases (green boxes) of the module 3 eigengene and the top-enriched GO BP terms and KEGG pathways in this module.

The first gene with the highest module membership in module 2 is *CAPN2* that is down-regulated in COPD cases (Fold-change = 0.8 and FDR = 7.3 ∙ 10^−7^) and encodes the large subunit of the ubiquitous enzyme, calpain 2. The calpains, calcium-activated neutral proteases, are nonlysosomal, intracellular cysteine proteases. It is worth emphasizing that smoking activates macrophages to produce a variety of inflammatory mediators including chemokines, reactive oxygen species (ROS), and proteases [King 2015]. Increasing evidence indicates that chronic inflammatory and immune responses play key roles in the development and progression of COPD [King 2015, Rovina 2013]. The chronic inflammatory process in COPD involves both innate (neutrophils, macrophages, eosinophils, mast cells, natural killer cells, T cells, innate lymphoid cells, and dendritic cells) and adaptive immunity (T and B lymphocytes) [Barnes 2016]. In particular, in patients with COPD, there is a characteristic pattern of lung inflammation with increased numbers of neutrophils, macrophages, and T and B lymphocytes. Consistently, we found that module 2 is more enriched in cell type-specific gene markers (i.e., immune gene signatures) known in literature [Nirmal 2018], with a total of 67 marker genes representative of six immune populations, all up-regulated in COPD cases (see Materials and Methods section, Figure 3 and Supplementary Table 2).

The first two genes with the highest membership in module 3 are *PRDX4* and *KCND3*, both up-regulated in COPD cases (Fold-change = 1.2 and FDR = 4.8 ∙ 10^−4^; Fold-change = 1.2 and FDR = 9.8 ∙ 10^−6^, respectively). *PRDX4* codes a protein that is an antioxidant enzyme belonging to the peroxiredoxin family. This protein is localized to the cytoplasm and has been found to play a regulatory role in the activation of the transcription factor NF-kappaB. Instead, the gene *KCND3* codes for potassium channel subfamily D member 3 and is located in a genome-wide significant COPD GWAS region, although the COPD-associated top SNP is located about 575 Kb from the transcription start site. The expression of *PRDX4* is strongly positively correlated in the COPD correlation network with *SERPINE2*, *CD79A*, and *POUF2AF1*, which were previously considered as putative interactors of genes at COPD GWAS loci [Morrow 2017]. The expression of *KCND3* is strongly positively correlated with two of them (*SERPINE2* and *CD79A*). Among KEGG pathways and GO Biological Processes enriched in module 3, we found annotations related to the regulation of the immune system and inflammatory response (see Materials and Methods section and Figure 3).

### Identification and characterization of switch genes

We classified nodes in the COPD correlation network using the date/party/fight-club hub classification system [Paci 2017], based on the Average Pearson Correlation Coefficients (APCCs) between the expression profiles of each hub and its nearest neighbors (see Materials and Methods section). Then, we assigned a topological role to each node based on their inter- and intra-cluster interactions and thus drew the heat cartography map for the COPD dataset, where party, date, and fight-club hubs were easily identified by red, orange, and blue coloring, respectively (Figure 4a and Supplementary Table 2). Through the heat cartography map, we are able to identify switch genes as a special subclass of fight-club hubs (APCC <0) characterized by having more links outside than inside their own cluster, while not being hubs in their own cluster (i.e., switch genes are fight-club hubs falling in the R4 region of the heat cartography map).

**Figure 4.**
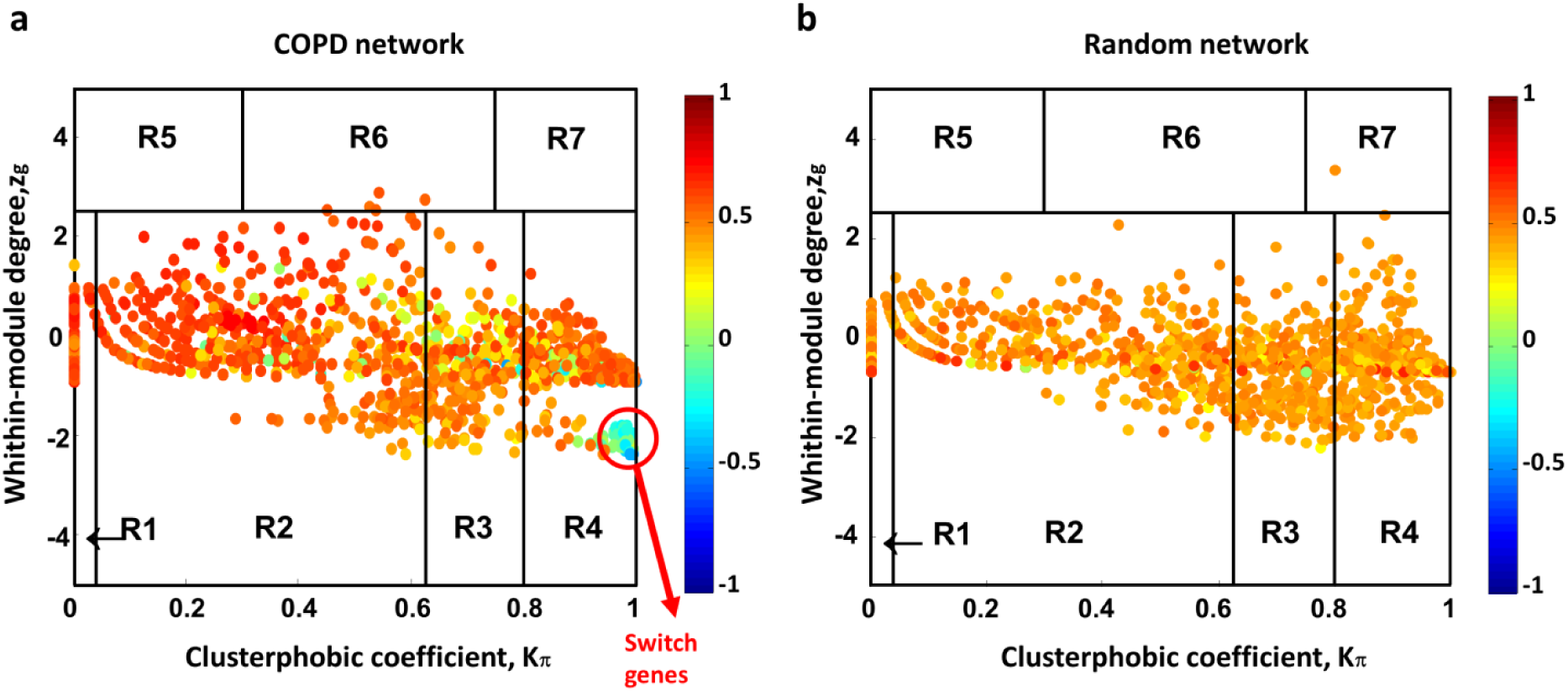
Identification of COPD switch genes. (a-b) Heat cartography maps of COPD and randomized network. Dots correspond to nodes in the networks. Each node is colored according to the value of the APCC between its expression profile and that of its nearest neighbors in the network. The randomized network was obtained by randomly shuffling the edges but preserving the degree of each node.

By drawing the heat cartography map for the nodes of the COPD randomized network (i.e., obtained by shuffling the edges but preserving the degree of each node), we observed a predominance of positive correlations and an absence of switch genes (Figure 4b). To assess statistical significance to this observation, we repeated this procedure 1000 times and we calculated the number of switch genes in each COPD randomized network. We found that the number of switch genes in each randomized network was always less than three, with a mean of 0.6 and a standard deviation of 0.8. Then, the number of 62 switch genes found in the original COPD correlation network (Figure 4a) was zscore-normalized and the p-value for the given z statistics was calculated; the p-value ~ 0 indicates that the observed heat cartography map in the COPD gene expression dataset (Figure 4a) is not a random event.

We found 62 switch genes in the COPD correlation network all resulting in gene up-regulation in COPD cases (Figure 5a and Supplementary Table 4). Most switch genes (74%) fall in module 3 (Figure 5b) and, mutually, module 3 is almost entirely populated by switch genes (73%), thus conferring to this module a well-characterized and defined signature as switch module. Note that *PRDX4* and *KCND3*, the first two genes with the highest membership in module 3, are also switch genes.

**Figure 5.**
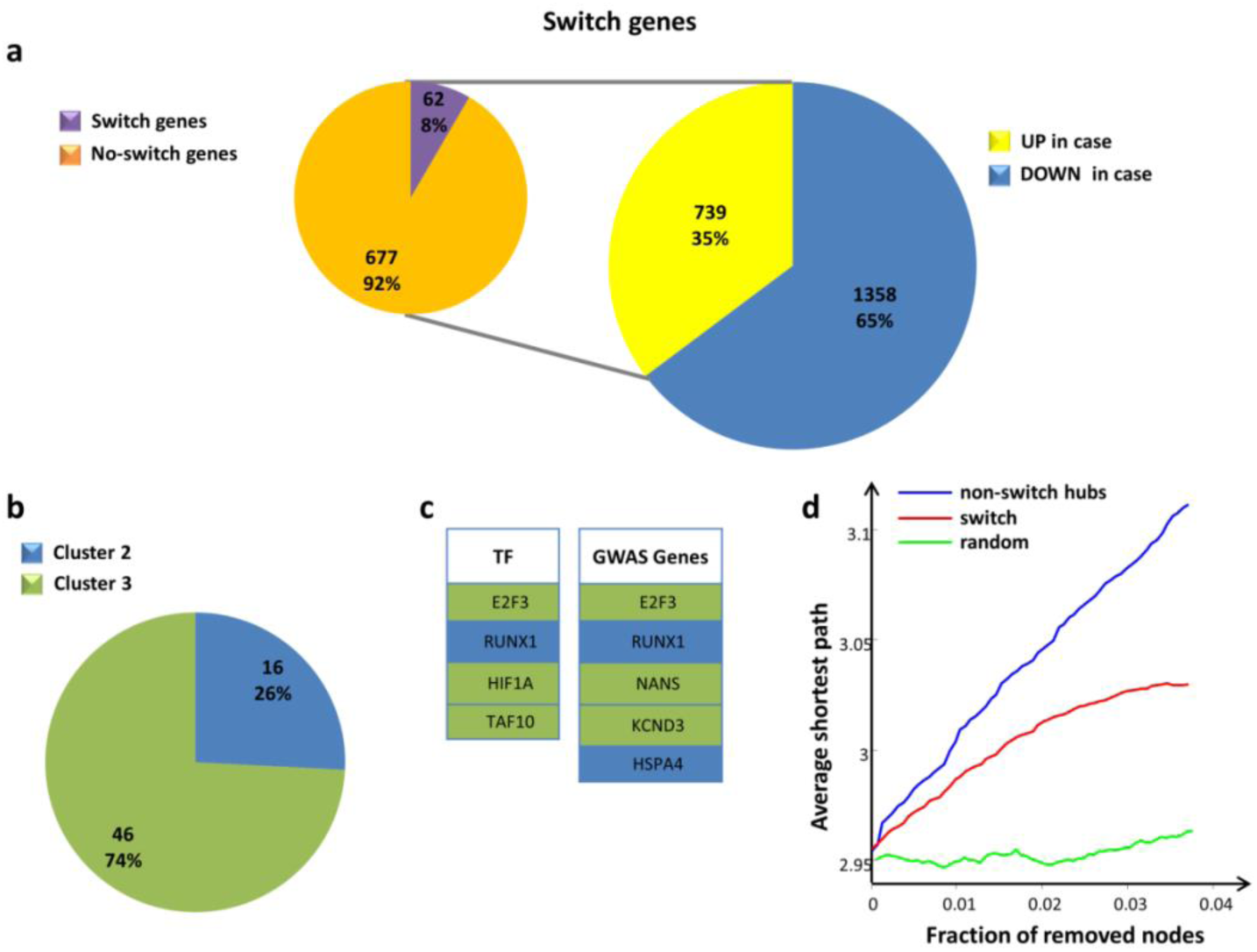
Characterization of COPD switch genes. (a) The larger pie chart [right] represents the percentages of DEGs that are up-/down-regulated in COPD cases in comparison to control subjects. The smaller pie chart [left] represents the percentages of switch genes among the up-regulated genes in COPD cases. (b) The pie chart represents the percentages of switch genes in each cluster. Note that switch genes are mostly included in cluster 3, and no switch genes were found in cluster 1. (c) Tables listing the switch genes that are transcription factors [left] and GWAS genes [right]. Switch genes are colored according to their associated cluster. (d) Robustness of the COPD correlation network. Blue curve corresponds to the cumulative deletion of non-switch hubs (i.e., the first 62 hubs that are not switch genes, sorted by decreasing degree); red curve corresponds to the cumulative deletion of the 62 switch genes, sorted by decreasing degree; the green curve corresponds to the cumulative deletion of randomly selected nodes. The x-axis represents the cumulative fraction of removed nodes with respect to the total number of 1655 network nodes (i.e., x-maximum is 62/1655 = 0.04), while the y-axis represents the average shortest path.

COPD switch genes are all protein-coding, among which 4 transcription factors - including *E2F3*, *HIF4*, *TAF10* of module 3 and *RUNX1* of module 2 - and five other genes located within previously identified genome-wide significant COPD GWAS loci [Sakornsakolpat 2019] (Figure 5c).

Functional annotation analysis of switch genes reveals that they are mainly involved in the regulation of several functionalities related to the immune and inflammatory response, mirroring the enrichment results obtained for module 3 (Supplementary Figure 3).

### Removal of switch genes

Scale-free networks display a surprising degree of tolerance against errors and the ability of nodes to communicate is unaffected even by very high node failure rates. For example, organisms grow and reproduce despite drastic environmental interventions and error tolerance is usually attributed to the robustness of the underlying metabolic network. However, this error tolerance comes at a high price in that these networks are extremely vulnerable to attacks, i.e., to the selection and removal of a few nodes that play a vital role in maintaining the network’s connectivity [Albert 2000].

We studied the tolerance of the COPD network against the removal of the 62 switch genes by comparing to the impact of the removal of the first 62 hubs that are not switch genes (called “non-switch hubs”). Both switch genes and non-switch hubs were sorted by decreasing degree and selected to be removed. Then, the effect on the average shortest path (where the shortest path between two nodes is the minimum number of edges connecting them, and the average shortest path is the mean of the shortest paths for all possible pairs of nodes in the network) of the cumulative node deletion is evaluated.

We found that the removal of switch genes produces a drastic increase of the average shortest path, mirroring the effect caused by the deletion of the first 62 non-switch hubs (Figure 5d). This means that switch genes play a vital role in maintaining the network’s connectivity while not being the primary hubs. In fact, the first 62 nodes sorted by decreasing degree include only two switch genes (Figure 6). On the contrary, the removal of 62 randomly selected nodes does not affect the integrity of the network (Figure 5d).

**Figure 6.**
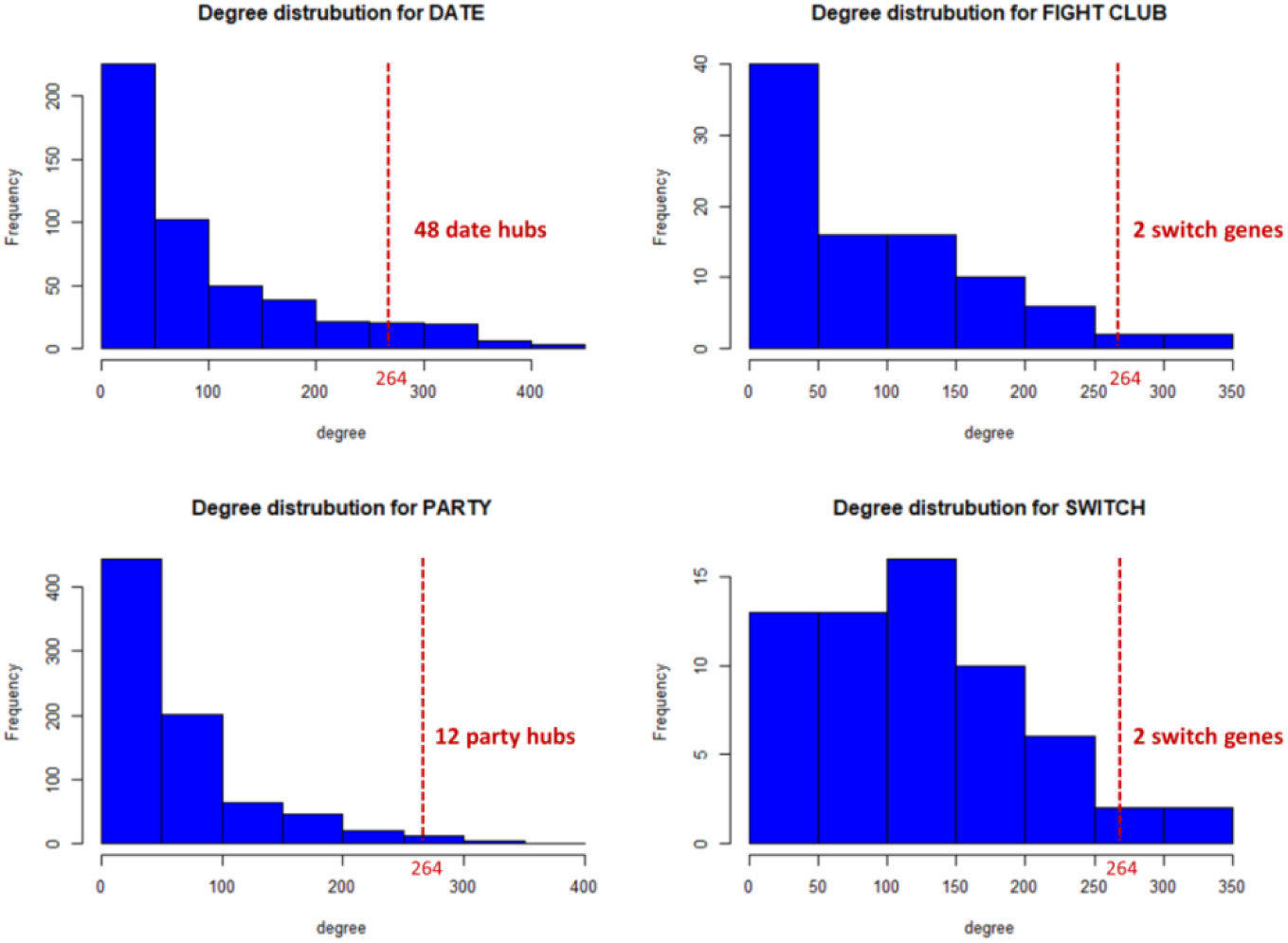
Degree distribution for each class of hubs. The dashed red lines correspond to the lowest degree (i.e., 264) of the first 62 (i.e., number of switch genes) nodes sorted by decreasing degree. For each class of hubs, the number of nodes that are included in the first 62 sorted nodes is reported.

### Validation of switch gene identification

To further assess the validity of the SWIM analysis in identifying disease genes and modules associated to COPD status, we applied the SWIM software on the GSE76925 dataset (test set), which contains microarray gene expression profiling of lung tissue samples from 111 COPD cases and 40 control smokers with normal lung function [Morrow 2017]. In this case, starting from 22631 genes, we obtained 887 significantly DEGs at 15% FDR, of which 493 (56%) were down-regulated in COPD cases, while the remaining 394 (44%) were up-regulated (Supplementary Table 1). To build the COPD correlation network, we selected a correlation threshold equal to 0.55 (corresponding to the 95^th^ percentile of the entire correlation distribution), which roughly guarantees to preserve the network integrity. A higher correlation threshold would cause a drastic drop in network connectivity.

The obtained COPD correlation network encompassed 667 nodes and 22595 edges, including 103 date hubs, 348 party hubs, and 71 fight-club hubs. From the COPD correlation network, SWIM extracted 61 switch genes all resulting in gene up-regulation in COPD cases (Supplementary Figure 4a, Supplementary Table 4). By studying the tolerance of the COPD correlation network against the removal of the 61 switch genes, similar to the other lung tissue gene expression dataset, we found that the removal of switch genes produces a drastic increase of the average shortest path, even overcoming the effect caused by the deletion of the first 61 non-switch hubs (Supplementary Figure 4b). This strongly supports the hypothesis of their putative key role in preserving the network’s connectivity, while not being the primary network hubs (the first 61 nodes sorted by decreasing degree do not include any switch gene).

Notably, the list of 61 switch genes includes *SPP1* that encodes adiponectin, which has been suggested as a protein biomarker for COPD [Suh 2018] and *TUFM*, which is probably the best candidate within a strong COPD GWAS region on chromosome 16 [Hobbs 2016].

The two analyzed datasets shared only one switch gene, i.e. *SSR4*, which appears in the top-ten switch genes of the training set (GSE47460 dataset). Previous analyses of gene expression differences in COPD have noted the challenges of finding consistent results across studies [Hobbs 2014]. There is marked heterogeneity in the development of COPD even among people with similar cigarette smoking histories, which is likely partially explained by genetic variation making the functional understanding of the disease a formidable challenge. Network-based approaches enable modeling of the complex molecular interactions involved in COPD pathogenesis aiding translational understanding of the complex mechanisms underlying the disease. These approaches start from the assumption that complex diseases are rarely a consequence of an abnormality in a single gene, but are likely influenced by a network of interacting genes and proteins where diseases can be identified with localized perturbation within a certain neighborhood [Barabási 2011]. The identification of these neighborhoods is therefore a prerequisite of a detailed investigation of a particular pathophenotype. In order to investigate the extent to which the two lists (S1, S2) of switch genes are in the immediate vicinity of each other in the Human Interactome (i.e., the cellular network of all physical molecular interactions), we used a network proximity measure and interactome database obtained from [Cheng 2018]:

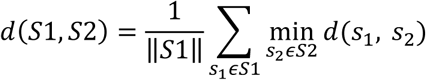

where the closest distance *d*(*S*1, *S*2) is the average shortest path length between switch genes *s*_1_ of the list *S*1 and the nearest switch gene *s*_2_ of the list *S*2 (Figure 7).

To evaluate the significance of the observed network proximity across the two lists of switch genes, we built a reference distance distribution corresponding to the expected distance between two randomly selected groups of proteins of the same size and degree distribution as the original two sets of switch genes in the human interactome. This procedure was repeated 5000 times (Figure 7).

**Figure 7.**
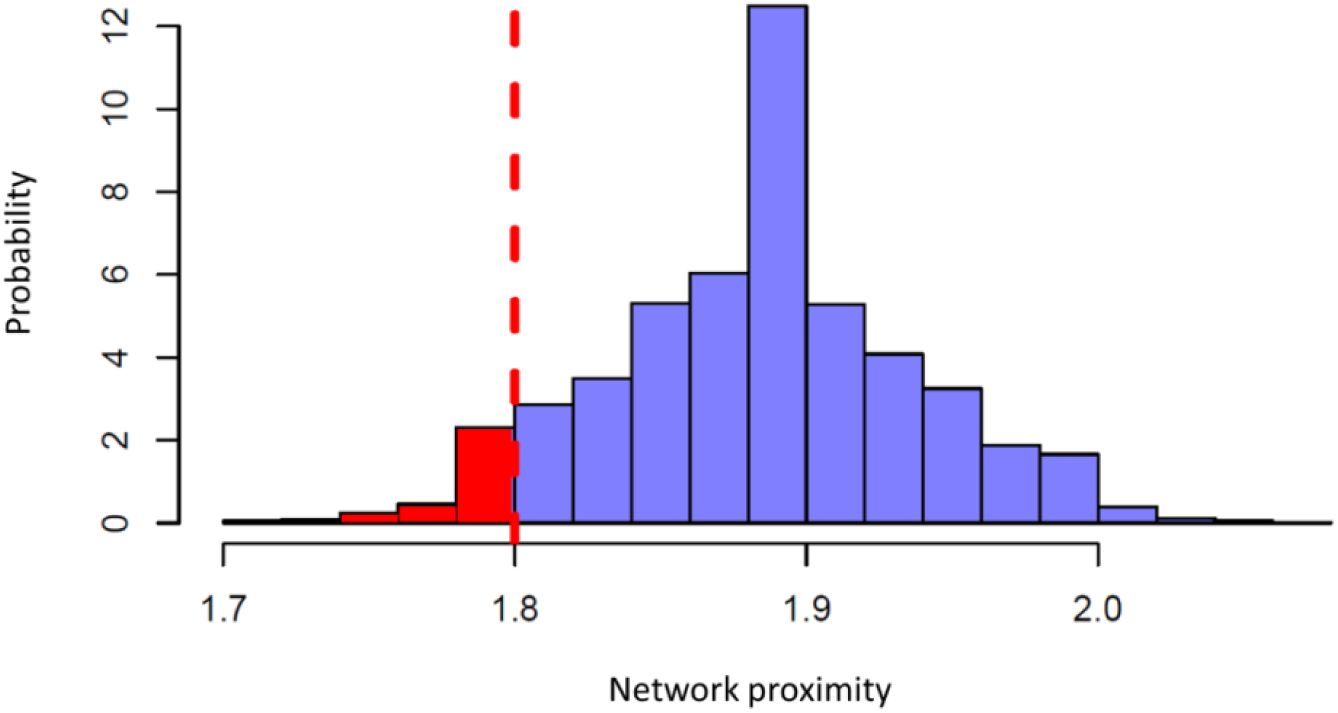
Probability distribution of the network proximity. The dashed red line corresponds to the observed network proximity measure (*d*=1.8) across the lists of switch genes in the two analyzed datasets. The red area represents the probability of observing the test statistic as small as that observed, corresponding to a p-value = 0.049, or smaller.

Then, the network proximity measure across the two lists of switch genes was zscore-normalized by using the mean and the standard deviation of the reference distance distribution. Subsequently, the p-value for the given z statistics was calculated. The obtained p-value < 0.05 indicates that the proximity in the human interactome of the two lists of switch genes is lower than the mean of the network distances between any two sets of randomly selected nodes of the same size and degree.

Interestingly, by studying the functional annotations of the two lists of switch genes, we found that they share COPD-related KEGG pathways (Table 1) and GO biological processes (Table 2), namely the NF-κB and toll-like receptor signaling pathways, regulation of immune and inflammatory response and key processes in cell development. Among switch genes involved in the NF-κB pathway, we found *SYK*, *BLNK*, and *LY96* in the training set (GSE47460 dataset) and *IRAK4* in the test set (GSE76925 dataset), all up-regulated in COPD cases. Thus, despite the apparent discrepancy in the lists of switch genes across the two datasets, the observed proximity in the human interactome, as well as the shared COPD related functionalities suggest they are working together in determining the COPD pathophenotype.

**Table 1.** Common KEGG functional annotations. Table showing the KEGG pathways shared between the two list of switch genes obtained from the training and test set (i.e., GSE47460 and GSE7925).

| KEGG pathways | switch GSE47460 | switch GSE76925 |
| --- | --- | --- |
| Antigen processing and presentation | *HSPA4* | *HSP90AA1*; *CD8A* |
| Th17 cell differentiation | *HIF1A*; *RUNX1* | *HSP90AA1* |
| Primary immunodeficiency | *BLNK* | *CD8A* |
| NF-kappa B signaling pathway | *SYK*; *BLNK*; *LY96* | *IRAK4* |
| Toll-like receptor signaling pathway | *LY96* | *SPP1*; *IRAK4* |
| NOD-like receptor signaling pathway | *IFI16*; *CYBA* | *HSP90AA1*; *IRAK4* |
| PI3K-Akt signaling pathway | *SYK*; *EIF4EBP1* | *HSP90AA1*; *SPP1* |
| Cellular senescence | *CHEK2*; *EIF4EBP1*; *E2F3* | *RAD50* |

**Table 2.** Common GO BP functional annotations. Table showing the GO Biological Processes shared between the two list of switch genes obtained from the training and test set (i.e., GSE47460 and GSE7925).

| GO Biological Process | Switch GSE47460 | Switch GSE76925 |
| --- | --- | --- |
| antigen processing and presentation of exogenous peptide antigen via MHC class I | *CYBA* | *PSMC2* |
| antigen receptor-mediated signaling pathway | *SYK* | *PSMC2* |
| cellular response to cytokine stimulus | *TIMP1;IL13RA2;HIF1A* | *HSP90AA1;IRAK4;EPRS* |
| cellular response to interleukin-1 | *HIF1A* | *PSMC2;IRAK4* |
| cytokine-mediated signaling pathway | *SYK;RPLP0;TIMP1;IL13RA2;HIF1A* | *HSP90AA1;PSMC2;IRAK4* |
| innate immune response activating cell surface receptor signaling pathway | *SYK* | *PSMC2* |
| positive regulation of T cell proliferation | *SYK* | *CD24* |
| regulation of cytokine-mediated signaling pathway | *SYK;RUNX1* | *CD24* |
| regulation of immune response | *SYK* | *SELL;CD8A* |
| regulation of interleukin-2 production | *RUNX1* | *NAV3* |
| negative regulation of interleukin-2 production | *TRIM27* | *NAV3* |
| neutrophil activation involved in immune response | *PRDX4;SYK;CYBA* | *HSP90AA1;SELL;COPB1;PSMC2;YPEL5;GYG1* |
| neutrophil degranulation | *PRDX4;CYBA* | *HSP90AA1;SELL;COPB1;PSMC2;YPEL5;GYG1* |
| neutrophil mediated immunity | *PRDX4;CYBA* | *HSP90AA1;SELL;COPB1;PSMC2;IRAK4;YPEL5;GYG1* |
| neutrophil migration | *SYK* | *IRAK4* |
| regulation of response to cytokine stimulus | *RUNX1* | *CD24* |
| positive regulation of I-kappaB kinase/NF-kappaB signaling | *NEK6* | *IRAK4* |
| regulation of I-kappaB kinase/NF-kappaB signaling | *NEK6* | *IRAK4* |
| toll-like receptor signaling pathway | *LY96* | *IRAK4* |
| MyD88-dependent toll-like receptor signaling pathway | *LY96* | *IRAK4* |
| cellular response to hypoxia | *HIF1A* | *PSMC2* |
| extracellular matrix disassembly | *TIMP1* | *SPP1* |
| extracellular matrix organization | *GREM1;CYP1B1;TIMP1* | *SPP1* |
| cellular response to DNA damage stimulus | *BLM;CHEK2* | *RAD50* |
| regulation of cellular response to stress | *NEK6* | *HSP90AA1* |
| negative regulation of cell adhesion mediated by integrin | *CYP1B1* | *PDE3B* |
| negative regulation of angiogenesis | *SERPINF1* | *PDE3B* |
| negative regulation of apoptotic process | *GREM1* | *TOX3* |
| regulation of cell differentiation | *GREM1;RUNX1* | *CD24* |
| regulation of cell migration | *CYP1B1* | *NAV3* |
| negative regulation of cell migration | *CYP1B1;TIMP1* | *NAV3* |
| negative regulation of cell motility | *CYP1B1* | *NAV3* |
| negative regulation of cell proliferation | *GREM1;CYP1B1;E2F3;NME1* | *CTCF* |
| regulation of cell proliferation | *MCTS1;GREM1;SYK;CYP1B1;TIMP1;NME1* | *CTCF* |
| regulation of intracellular signal transduction | *ARHGAP22;BLM;CHEK2;TAF10* | *RAD50;CD24* |
| regulation of signal transduction | *TIMP1;RUNX1* | *CD24* |
| regulation of signal transduction by p53 class mediator | *BLM;CHEK2;TAF10* | *RAD50* |
| regulation of stem cell differentiation | *RUNX1* | *PSMC2* |
| negative regulation of Wnt signaling pathway | *GREM1* | *PSMC2* |

### Switch genes interacting with genes at COPD GWAS loci and with SERPINE2, CD79A and POUF2AF1

Looking at the COPD GWAS regional genes that are nearest network neighbors of switch genes in the training set (GSE47460 dataset), we found that switch genes are highly correlated and anti-correlated with 24 and 36 GWAS genes, respectively (Supplementary Table 5). Since a liberal region around 82 genomic loci associated with COPD was included (about +/− 1 Mb), approximately 5% of the genome was encompassed by those COPD GWAS regions. Thus, in order to check if the number of GWAS genes included in the positive/negative nearest neighbors of COPD switch genes (i.e., 24 and 36 GWAS genes, respectively) is more than expected by chance, the nearest neighbors of COPD switch genes were randomly shuffled 1000 times preserving the degree of each switch gene and the interaction weights. Then, the original values (non-random values) of GWAS positive and negative nearest neighbors were z-score-normalized and the p-values for the given z statistics were calculated, that are 6.63 × 10^−76^ and 3.93 × 10^−205^, respectively. This suggests that the observed number of GWAS genes included in the positive/negative nearest neighbors of COPD switch genes (i.e., 24 and 36 GWAS genes, respectively) is not a random event.

Interestingly, the list of 36 GWAS genes that are negative nearest neighbors of switch genes encompasses *AGER* and *EMP2*, which negatively correlate with 26 and 33 switch genes, respectively, of which 24 switch genes are in common (Figure 8 left and Supplementary Table 5). Note that this signature in mainly due to the switch genes falling in module 3. In fact, *AGER* and *EMP2* negatively correlates in module 3 with 25 and 30 switch genes, respectively, of which 23 switch genes in common, including *KCND3*, *SSR4*, *LY96*, and *TIMP1*. Overall, the 70% of the negative interactors of *AGER* and/or *EMP2* in module 3 are switch genes.

**Figure 8.**
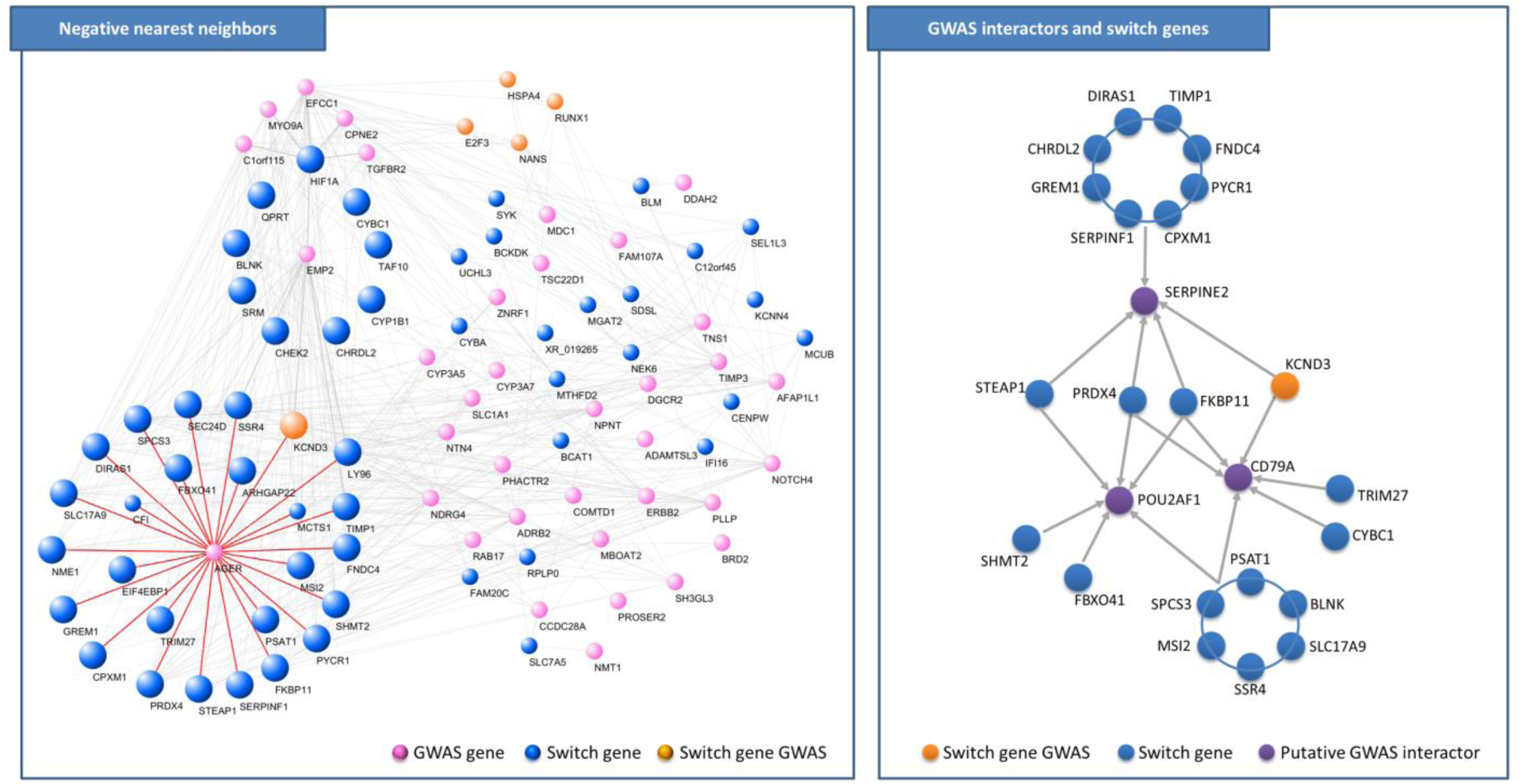
Switch genes interactions. [LEFT] Networks of switch genes negatively correlated with GWAS genes. Pink nodes correspond to GWAS genes, blue nodes correspond to switch genes, orange nodes correspond to switch genes that are also GWAS genes, larger size nodes correspond to negative nearest neighbor of *EMP2*. The interactions of *AGER* with its nearest neighbors are highlighted in red. [RIGHT] Sketched network of correlations among switch genes and *SERPINE2*, *CD79A*, and *POUF2AF1*.

Looking at genes that have been previously considered as putative interactors of genes at COPD GWAS loci [Morrow 2017], we found that 22 switch genes are strongly positively correlated with *SERPINE2*, or with *CD79A*, or with *POUF2AF1* (Figure 8 right). Among them, we found *PRDX4* and *FKBP11* that are positively correlated with all three GWAS interactors *POUF2AF1*, *SERPINE2*, and *CD79A*.

## Discussion

We analyzed lung tissue gene expression data from two well-characterized COPD case-control populations to study the differences between lung samples from normal subjects (represented by smokers with normal spirometry) and COPD cases. We used one dataset as a “training set” to perform the analysis and the other dataset as a “test set” to validate the results. We built a COPD correlation network, and we exploited the module-centric approach to identify putative COPD molecular determinants that we called “switch genes”.

COPD is characterized by lung inflammation that persists after smoking cessation. This inflammation is heterogeneous, but the key inflammatory cell types involved are macrophages, neutrophils, T and B cells. Macrophages play a key role in orchestrating chronic inflammation in patients with COPD and are found in markedly increased numbers in both the airways and lung parenchyma. Neutrophils provide powerful proteases and are most prominent in acute exacerbations in the lung airways [Tuder 2012]. The role of T and B cells in COPD pathogenesis has growing support from the basic science studies of COPD [Hogg 2004, Seys 2015, Polverino 2016, Morrow 2017]. In fact, recent evidence shows a strong correlation between increases in the number and size of B cell-rich lymphoid follicles and COPD severity, as well as a strong correlation between T cell numbers and the amount of alveolar destruction and severity of airflow obstruction [Barnes 2016, Polverino 2016]. Finally, B cell products (autoantibodies) were found in COPD blood and lung samples [Polverino 2016].

Multiple aspects of innate and adaptive immune functions are regulated by the transcription factor NF-κB that serves as a pivotal mediator of inflammatory responses. NF-κB induces the expression of various pro-inflammatory genes and plays a critical role in regulating the survival, activation and differentiation of innate immune cells and inflammatory T cells [Liu 2017]. Consequently, deregulated NF-κB activation contributes to the pathogenic processes of various inflammatory diseases. Recently, various therapeutic strategies that target the NF-κB signaling pathway have been considered for treatment of inflammatory diseases, such as asthma and COPD [Schuliga 2015]. Among the causes responsible for the activation of this pathway, there is the binding of the advanced glycosylation end-products (AGEs) to *RAGE*, the protein encoded by the gene *AGER* that is one of the most well-known candidate genes located in a significant COPD GWAS region. *RAGE* is a membrane receptor, but also has soluble forms (*sRAGE*) generated mainly by alternative splicing mechanism of the *AGER* gene. Reduced *sRAGE* levels are associated with heightened inflammation in various chronic conditions, and they are also associated with increased emphysema and COPD status [Kim 2015]. sRAGE is one of the most promising biomarkers for emphysema [Yonchuk 2015].

In addition to inflammation, our analyses highlighted hypoxia-related pathways. Inflammation shares an interdependent relationship with hypoxia. In fact, oxygen passes from the lung tissue to the blood via the lung alveoli. COPD damages the lungs, and if they get seriously damaged, hypoxia may occur since the blood does not deliver enough oxygen to the alveoli in the lungs. Patients with inflammatory diseases experience elevated levels of hypoxia-inducible factors (*HIF*), a transcription factor that is stabilized during conditions of hypoxia, and the activation of *HIF1* signaling pathway has been shown to correlate with a decrease of lung function, reduced quality of life and progression of COPD [Fu 2018, Rong 2018, Lo 2018]. Thus, while hypoxia can elicit tissue inflammation, inflammatory disease states are frequently characterized by tissue hypoxia, supporting the hypothesis that hypoxia and inflammation are two sides of the same coin [Bartels 2013]. Besides inducing inflammation, hypoxia causes also the disappearance of caveolae in the epididymal adipose tissue and inhibits the expression of *CAVIN1* through *HIF1* [Regazzetti 2015]. Caveolae dysfunction is implicated in several pathologies, such as muscular dystrophies and pulmonary hypertension in chronic obstructive pulmonary disease [Regazzetti 2015, Huber 2009].

Consistent with all these observations, we found that the COPD correlation network built by SWIM software consists of three well-characterized modules: one populated by switch genes all up-regulated in COPD cases and related to the regulation of immune and inflammatory response; one populated by well-recognized immune signature genes all up-regulated in COPD cases; and one where the GWAS gene *AGER* and *CAVIN1* are the most representative module genes, both down-regulated in COPD cases. Interestingly, 70% of the negative interactors of *AGER* are switch genes.

Among switch genes involved in NF-κB signaling pathway, we found *LY96*, *BLNK*, and *SYK* (Supplementary Figure 3). In particular: *LY96* codes a protein which is associated with toll-like receptor 4 on the cell surface and confers responsiveness to lipopolysaccharide (LPS), thus providing a link between the receptor and LPS signaling; *BLNK* codes for an adaptor protein that plays a crucial role in B cell development and activation since it bridges B cell receptor-associated kinase activation with downstream signaling pathways; *SYK* encodes a member of the family of non-receptor type Tyr protein kinases that is involved in coupling activated immunoreceptors to downstream signaling events mediating diverse cellular responses, like proliferation, differentiation, and phagocytosis. Among switch genes linked to the NF-κB signaling pathway, we found *PRDX4* that is associated with neutrophil activation and degranulation and with I-kappaB phosphorylation that is an important step in the NF-κB activation. Moreover, we found that the gene expression of *PRDX4* strongly correlates with the activation of known COPD GWAS interactors *SERPINE2*, *CD79A*, and *POUF2AF1* [Cha 2009, Groneberg 2004, Mbebi 1999, Ladjemi 2015, Teitell 2003, Zhou 2016, Farner 2016].

Among switch genes involved in the inflammatory response and hypoxia, sharing an interdependent relationship [Bartels 2013], we found *TIMP1* and the transcription factor *HIF1A* (Supplementary Figure 3). *TIMP1* is the tissue inhibitor of metalloproteinase-1 related to airway hyperresponsiveness (AHR) in smokers [Lo 2018]. AHR is associated with airway inflammation and more rapid decline in lung function and is a predictor of future risk of COPD among smokers [Lo 2018]. *HIF1A* encodes the alpha subunit of hypoxia-inducible factor-1 (*HIF1*) and has been shown to be an essential regulator of the response to hypoxia. *HIF1A* is known to mediate oxygen delivery through its effects on vascular remodeling and angiogenesis, and oxygen utilization through involvement in redox homeostasis and glucose metabolism [Shimoda 2011]. Recent data have also suggested that *HIF1A* plays a major role in COPD, indicating that its high expression may be associated with decreased lung function and reduced quality of life, contributing to disease progression [Rong 2018, Fu 2018].

Among switch genes related to the regulation of immune response, we found the gene *CYBA* associated with the nucleotide-binding oligomerization domain-like (NOD-like) receptor signaling pathway. NOD-like receptors represent a group of key sensors for microbes and damage in the lung and might also indirectly regulate immune responses. Thus, they play a key role in various infectious as well as acute and chronic sterile inflammatory diseases, such as pneumonia and COPD [Chaput 2013]. *CYBA* codes for light chain (alpha subunit) of the cytochrome b protein, which has been proposed as a primary component of the microbicidal oxidase system of phagocytes and shows selective cytoplasmic expression in immune cells.

Furthermore, we found also switch genes involved in other mechanisms beyond chronic inflammation that are implicated in the development and the progression of the COPD, such as cellular senescence, apoptosis, and oxidative stress (Supplementary Figure 3). Oxidative stress is a critical feature in patients with COPD and drives accelerated aging though activation of phosphoinositide 3-kinase (PI3K) and reduction in sirtuin-1 levels, which leads to cellular senescence and release of inflammatory proteins, which further increase oxidative stress [Rovina 2013, Barnase 2016].

Finally, we used the test set of gene expression samples to validate the results obtained with the training set. Interestingly, by studying the functional annotations of the two lists of switch genes, we found that they share COPD-related KEGG pathways and GO biological processes, namely the NF-κB and toll-like receptor signaling pathways, regulation of immune and inflammatory response and key processes in cell development.

### Limitations and future directions

In this study, we have collected a number of clues, ranging from global to local properties, from purely computational to more biological ones, aiming to draw a sketch of putative underlying mechanisms that could lead to a large-scale transition towards the occurrence of COPD. It is worth noting that the computational approach used in this study is based on correlations that are just “associations” and do not imply necessarily “causal” relationships. Nevertheless, adding further computational and biological information allowed us to zoom-in from global properties (i.e., power laws, fight club hubs, switch genes, etc.) to a small pool of genes that could give the promise for a better understanding of the molecular mechanisms underlying the onset of COPD.

Moreover, SWIM constructs hard-thresholded networks (or binary networks) in order to remove meaningless relationships and thus focus on significant associations between highly correlated nodes. The hard-thresholding approach creates binary networks where sub-threshold inter-node correlations are suppressed (edge values set to 0), and supra-threshold correlations are compressed (edge values set to 1). This approach could in principle lead to a loss of information since small differences in correlation strength, or in the chosen threshold, can result in edges being present or absent in the resulting network. However, this limitation is partially overcome by using more stringent thresholds, thus maximizing the contributions from the strongest correlations and emphasizing the network characteristics exhibited more strongly in the extreme (positive and negative) tails of the correlation distribution.

An efficient solution to the hard-thresholding problem is to build soft-thresholded networks where thresholding is replaced with a continuous mapping of correlation values into edge weights, which has the effect of suppressing rather than removing weaker connections. In the future we hope to improve the SWIM functionalities by proposing different types of “soft” adjacency functions, like a sigmoid or logistic function.

Other limitations of our analysis include the relatively small size of the gene expression datasets and the heterogeneous nature of lung tissue samples. Increasing samples size and using data from specific cell types in future studies will likely improve the identification of COPD molecular determinants.

## Conclusions

Our findings demonstrate that switch genes play an active role in inflammatory responses and regulating the immune environment in COPD. Modulating the function of switch genes may be an important mechanism to dampen the hypoxia-promoting inflammatory response and may lead to an improved understanding of COPD pathogenesis.

The majority of the genes highlighted through the SWIM methodology would not have been identified using a traditional GWAS approach. This observation demonstrates how SWIM can aid the identification and the prioritization of novel diagnostic markers or therapeutic candidate genes involved in the etiology of COPD.

## Materials and Methods

### Datasets

#### GSE47460 dataset

The first dataset analyzed for the present study is available through the GEO public repository at accession number GSE47460 published on May 30, 2013 [Peng 2016, Anathy 2018]. This dataset includes microarray gene expression profiling obtained from total RNA extracted from whole lung homogenates from subjects undergoing thoracic surgery for clinical indications. These subjects were diagnosed as being controls or having interstitial lung disease (ILD) or chronic obstructive pulmonary disease (COPD). All samples are from the Lung Tissue Research Consortium (LTRC) and derived from two array platforms with a total of 582 samples: 255 have ILD, 219 have COPD, 108 are controls.

In order to compare COPD cases versus control, the gene expression data GSE47460 was analyzed as follows: ILD samples were removed from the two array platforms and then each array-type was Robust Multi-array Average (RMA)-normalized separately [Irizarry 2003]. Then, to “stitch together” the data from the two arrays, we first matched genes based on probe-ids (the arrays were quite similar and had many overlapping probes). However, some genes were never measured by the same probe (e.g., IREB2). Therefore, we next matched any remaining genes based on shared gene-id. After creating a single-merged dataset with both array types together, we treated the array-types as “batches” and ran Combat function from R/Bioconductor package SVA to correct for array-specific effects. The probe-sets were mapped to official gene symbols by using BioMart – Ensembl tool (https://www.ensembl.org/).

#### GSE76925 dataset

The second dataset analyzed for the present study is available through the GEO public repository at accession number GSE76925 published on Mar 29, 2017 [Morrow 2017]. This dataset collects microarray gene expression profiling of lung or airway tissues from subjects with chronic obstructive pulmonary disease (COPD) by using HumanHT-12 BeadChips (Illumina, San Diego, CA). A total of 111 COPD cases and 40 control smokers with normal lung function were collected; all subjects were ex-smokers. The probe-sets were mapped to official gene symbols by using the platform GPL10558 (Illumina HumanHT-12 V4.0 expression beadchip) available from GEO repository. Multiple probe measurements of a given gene were collapsed into a single gene measurement by considering the mean.

#### Human protein–protein interactome

The human protein–protein interactome was downloaded from the Supplementary Data of [Cheng 2018]. The authors of [Cheng 2018] assembled 15 commonly used databases with multiple types of experimental evidence (including kinase-substrate interactions from literature-derived low-throughput and high-throughput experiments; binary PPIs from three-dimensional protein structures; signaling networks from literature-derived low-throughput experiments; literature-curated PPIs identified by affinity purification followed by mass spectrometry, Y2H, and/ or literature-derived low-throughput experiments) and their inhouse systematic human protein–protein interactome. This updated version of the human interactome is composed of 217,160 protein–protein interactions (edges or links) connecting 15,970 unique proteins (nodes).

### COPD GWAS genes

COPD genetic risk loci were extracted from [Sakornsakolpat 2019], where the authors performed a genome-wide association study (GWAS) for a total of 257,811 individuals (i.e., 35,735 cases and 222,076 controls) from 25 different studies, including studies from International COPD Genetics Consortium [Hobbs 2017] and UK Biobank [Sudlow 2015], that collected genetic and phenotypic data with lung function and cigarette smoking assessment. In particular, the authors of [Sakornsakolpat 2019] tested the association of COPD and 6,224,355 variants, identifying 82 loci associated at genome-wide significance (p-value < 5 ∙ 10^−8^). Genetic loci were defined in [Sakornsakolpat 2019] by using a 2 Mb window (+/− 1 Mb) around a lead variant (top SNP).

### Differentially expressed genes

To compute the differentially expressed genes (DEGs), we used R statistical software (v 3.4.4) and the package limma (Figure 14). For each dataset, we fitted a linear regression model (Table 3) to the expression values of each gene (EXP) in order to detect the association with the variable of interest representing the case/control condition (COPD). Microarray batch effects were addressed by using age, sex, and smoking status (i.e., current, ever, never) for GSE47460 dataset and age, sex, race and pack-years of smoking for GSE76925 dataset as clinical phenotypes. For both datasets, two surrogate variables (obtained via the R/Bioconductor package SVA) were added as further covariates in the linear models (Table 3). The linear models were fitted by using least squares regression. Then, an empirical Bayes shrinkage method was used by the package limma to obtain a moderated t-test statistic and its p-value. Adjustment for multiple testing were controlled for false discovery rate (FDR) method [Benjamini-Hochberg 1995].

**Table 3.** Linear regression models for association with the variable of interest. In this table the linear regression models used to fit each dataset were reported, where EXP refers to the gene expression data, and COPD refers to the variable of interest (i.e., case/control condition). Smoker status of GSE47460 dataset corresponds to: current, ever, or never.

| Dataset | Reference | GEO Accession | Linear Model |
| --- | --- | --- | --- |
| training set | Peng 2016 ,<br>Anathy 2018 | GSE47460 | EXP ~ COPD + age + sex + smoker<br>status<br>+ 2 surrogate_variables |
| test set | Morrow et al<br>2017 | GSE76925 | EXP ~ COPD + age + sex + race + pack<br>years + 2 surrogate_variables |

### SWIM software

In order to identify switch genes associated with the transition between control smokers and COPD cases, we run SWIM (Figure 9), a software for gene co-expression network mining developed in MATLAB with a user-friendly Graphical User Interphase (GUI) and freely downloadable [Paci 2017].

**Figure 9.**
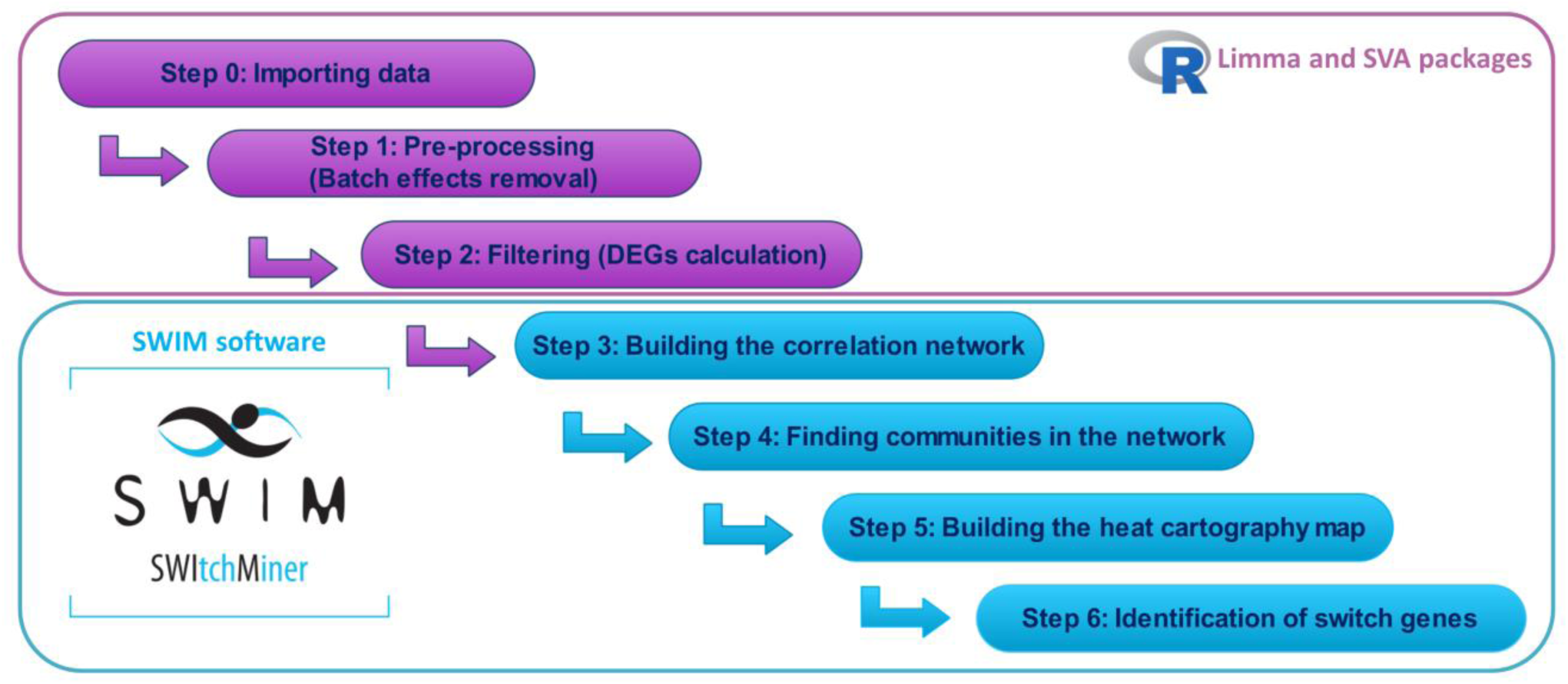
Flowchart of gene expression analysis.

SWIM builds a correlation network of differentially expressed genes. Usually, a network corresponds to an adjacency matrix *A* = [*a*_*i,j*_] that encodes the connection strength between each pair of nodes. In unweighted and undirected networks, *a*_*i,j*_ is equal to 1 if nodes *i* and *j* are connected and 0 otherwise. In particular, in unweighted and undirected gene correlation networks, *a*_*i,j*_ is equal to 1 if the expression profiles for nodes (i.e., genes) *i* and *j* are significantly associated across samples. In order to select significant associations, SWIM uses the absolute value of the Pearson correlation coefficient as similarity index. In other words, *a*_*i,j*_ is equal to 1 if the absolute value of the Pearson correlation coefficient between the expression profiles of nodes *i* and *j* is greater than a selected significance threshold. For the COPD correlation network, we set the correlation threshold equal to 0.57, which corresponded to the 98^th^ percentile of the entire correlation distribution. This choice for the correlation threshold stems from two selection criteria (Supplementary Figure 1). The first criterion is motivated by the observation that most biological networks display a scale-free distribution of node degree. Therefore, the network obtained based on the selected correlation threshold should approximate this topology. Scale-free networks are extremely heterogeneous, and their topology is dominated by a few highly connected nodes (hubs), which link the rest of the less connected nodes. Indeed, the defining property of scale-free networks is that the probability that a node is connected with *k* other nodes (i.e., the degree distribution *P*(*k*) of a network) decays as a power law *P*(*k*)~*k*^−*α*^. Many biological networks have been shown to be scale-free networks [Jeong 2000, Han 2004, Carter 2004]. For evaluating whether the COPD gene expression correlation network exhibits a scale-free topology, we calculated the square of the correlation between log (*P*(*k*)) and log (*k*), i.e. the index R-squared, as a function of the Pearson correlation (Supplementary Figure 1a). Since it is biologically implausible that a network contains more hub genes than non-hub genes, we multiply R-squared with −1 if the slope *α* of the regression line between log (*P*(*k*)) and log (*k*) is positive and thus we obtain a signed version of this index. If the R-squared approaches 1, then there is a straight-line relationship between log (*P*(*k*)) and log(*k*) and a scale-free topology is reached. These considerations motivated us to choose a correlation threshold that can lead to a network satisfying scale-free topology at least approximately, e.g. signed R-squared > 0.8 [Zhang 2005]. The second criterion relies on choosing a threshold that should reflect an appropriate balance between the number of edges and the number of connected components of the network: the number of edges should be as small as possible in order to have a manageable network (pointing towards a higher threshold) and the number of connected components should be as small as possible in order to preserve the integrity of the network (pointing towards a smaller threshold) (Supplementary Figure 1b).

Next, SWIM searches for specific topological properties of the correlation network using the date/party/fight-club hub classification system, based on the Average Pearson Correlation Coefficients (APCCs) between the expression profiles of each hub (i.e., node with degree greater than 5 [Han 2004]) and its nearest neighbors. Given a node *i* and its *n*_*i*_ first nearest neighbors, the APCC value is:

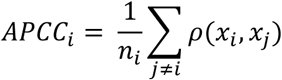

where ρ(*x*_*i*_, *x*_*j*_) is the Pearson correlation between the expression profiles of node *i* and its *j*-th nearest neighbor. The authors in [Paci 2017], defined: date hubs as hubs with APCC < 0.5 (i.e., low co-expression with their partners); party hubs as hubs with APCC ≥ 0.5 (i.e., high co-expression with their partners); and fight-club hubs as hubs with negative APCC values (i.e., inversely correlated with their partners). In the COPD network, SWIM found 92 fight-club hubs, 489 date hubs, and 795 party hubs.

SWIM then identifies communities in the network by means of the k-means clustering algorithm, employing Sum of Squared Errors (SSE) values to determine the appropriate number of clusters, and assigns a role to each node by using the Guimera-Amaral approach [Han 2004], based on the inter and intra-clusters interactions of each node quantified by the computation of two statistics: the within-module degree *z*_*g*_ and the clusterphobic coefficient *K*_*π*_. The two parameters are defined as:

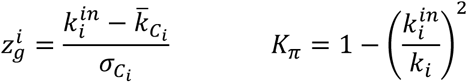

where 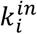 is the number of edges of node *i* to other nodes in its module *C*_*i*_, *k*_*i*_ is the total degree (i.e., number of edges emanating from a node) of node *i*, 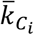 and 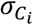 are the average and standard deviation of the total degree distribution of the nodes in the module *C*_*i*_. Roughly, the clusterphobic coefficient measures the “fear” of being confined in a cluster, in analogy with the claustrophobic disorder and the global within-module degree *z*_*g*_ measures how “well-connected” each node is to other nodes in its own community. According to *K*_*π*_ and *z*_*g*_ values, the plane is divided into seven regions (R1-R7), each defining a specific node role. High *z*_*g*_ values correspond to nodes that are hubs within their module (local hubs), while high values of *K*_*π*_ identify nodes that interact mainly outside their community, i.e., having much more external than internal links. SWIM colored each node in the plane identified by *z*_*g*_ and *K*_*π*_ according to its APCC value, thus defining a heat cartography map.

Finally, SWIM extracts a select set of genes, named switch genes, as a special subclass of fight-club hubs falling in the R4 region. In particular, switch genes have to satisfy the following topological and expression features:

1. Not being a hub in their own cluster (*z*_*g*_ < 2.5)
2. Having many links outside their own cluster (*K*_*π*_ > 0.8)
3. Having a negative average weight of their incident links (APCC < 0)

### Immune response signatures

Immune cell-related genes were obtained from [Nirmal 2018], where the authors identified 569 marker genes representative of seven immune populations: B cells (37 genes), plasma cells (14 genes), monocytes (37 genes), macrophages (78 genes), neutrophils (47 genes), NK cells (20 genes), T cells (85 genes). The authors of [Nirmal 2018] validated the data-driven definition of each immune signature by association of known markers with the specific gene signatures, e.g., CD3D and CD3E (T cells), CD19, CD22, and CD79 (B cells), CD14 (monocytes), CD68 and CD163 (macrophages), KIR family (NK cells) and immunoglobulin family members (plasma cells).

### Functional enrichment analysis

The associations between selected genes and functional annotations such as KEGG pathways [Kanehisa 2015] and GO terms [Ashburner 2000] were obtained by using Enrichr [Kuleshov 2016] web tool. P-values were adjusted with the Benjamini-Hochberg method and a threshold equal to 0.05 was set to identify functional annotations significantly enriched amongst the selected gene lists.

## Acknowledgments

This work was financial supported by NIH grants P01 HL114501, R01 HL137927, and R01 HL147148.

## Author contributions

PP, ES, and LF concept and design. PP, GF, FC, VL analysis of data. All authors contributed to interpretation of data, review and approval the final manuscript.

---

**Supplementary Figure 1.**
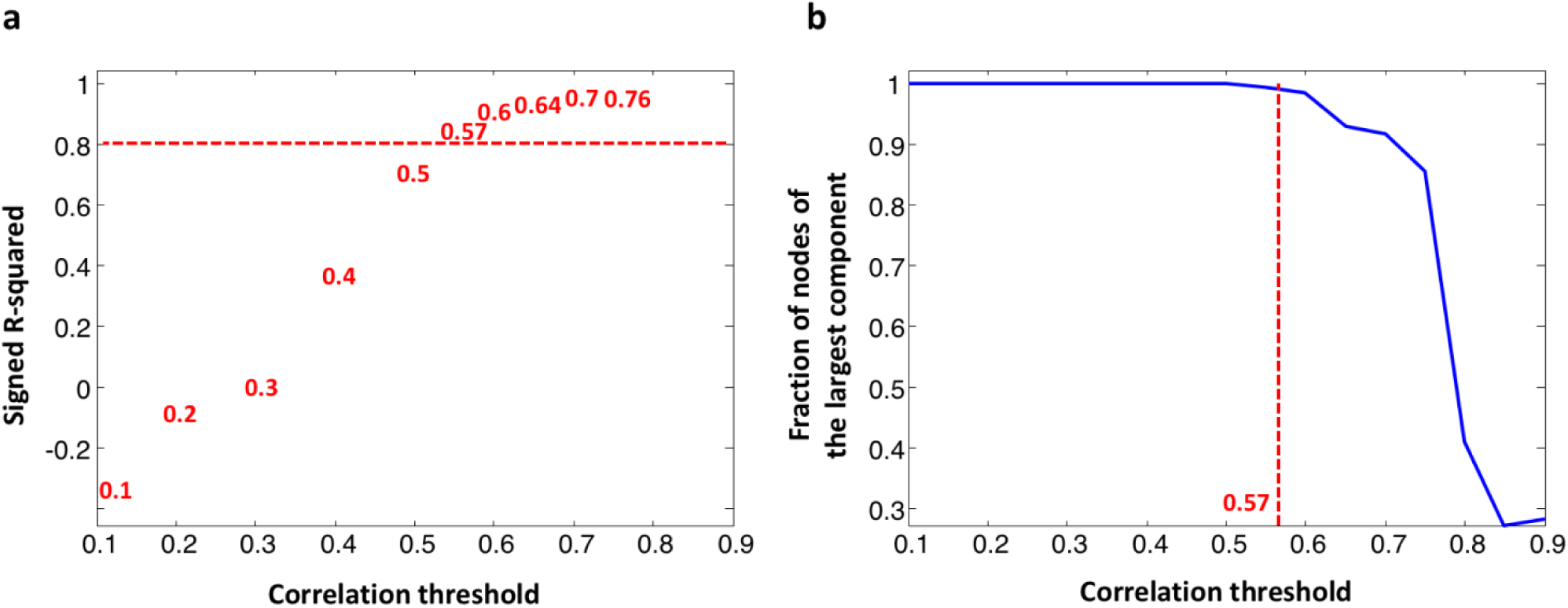
Correlation threshold for COPD network. (a) Analysis of network topology for various correlation thresholds. The x-axis represents the Pearson correlation threshold varying in the chosen range, while the y-axis represents the scale-free fit index (signed R-squared). The horizontal red line represents the smallest value of signed R-squared (0.8) such that an approximate scale-free topology is reached. (b) Connectivity of the COPD correlation network for various correlation thresholds. The x-axis represents the Pearson correlation threshold varying in the chosen range, while the y-axis represents the fraction of nodes populating the largest component. The dashed red line corresponds to the selected threshold (*ρ* = 0.57 or 98^th^ percentile). Note that y=1 means that all nodes fall in the largest component and thus the network is fully connected; otherwise more components exist.

**Supplementary Figure 2.**
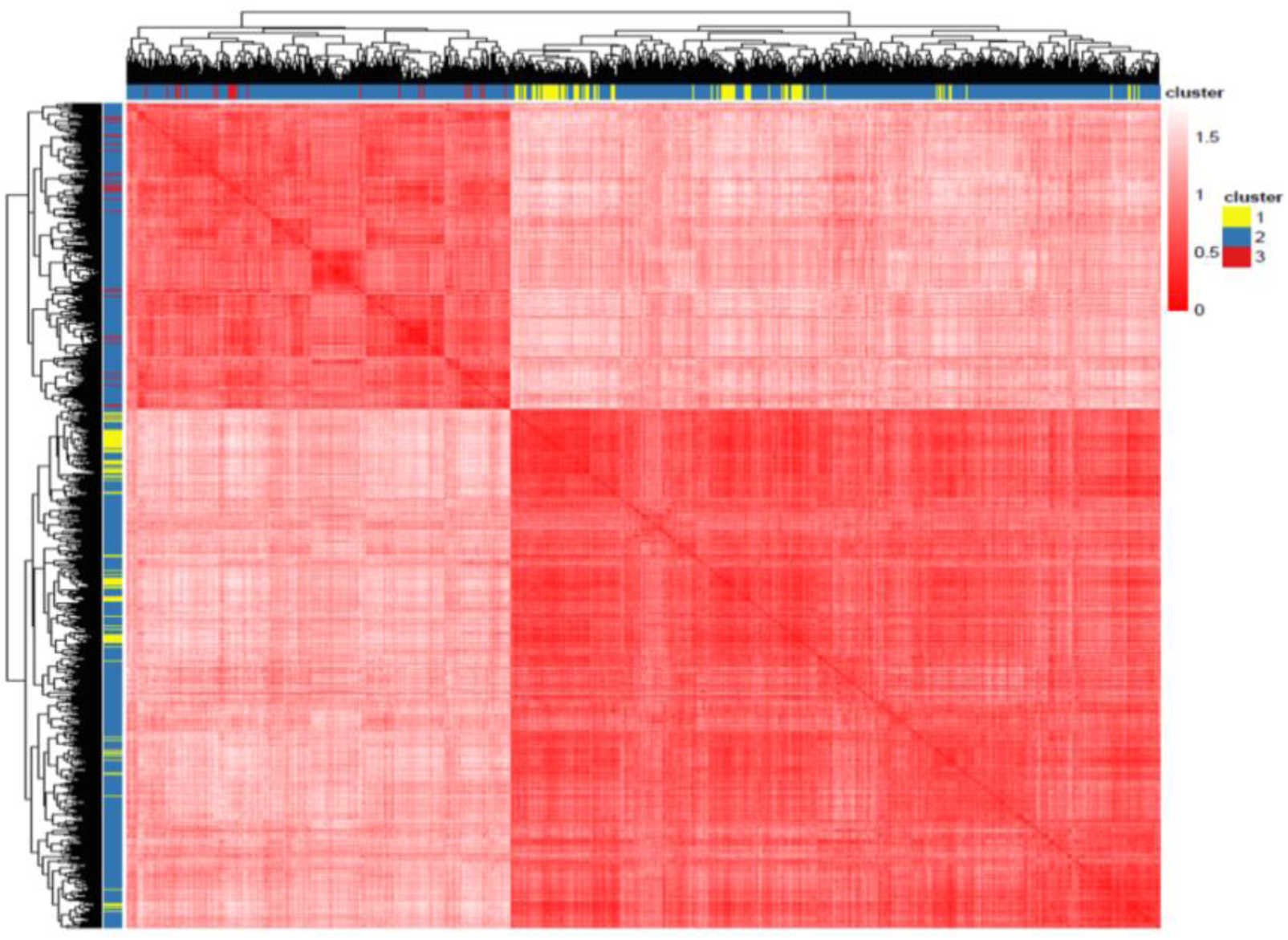
Hierarchical clustering for COPD network. Heat map of the values of the correlation-based dissimilarity where red/white colors indicate high/low values of dissimilarity. Here, the network nodes (rows and columns) have been clustered according to hierarchical clustering with complete agglomeration method and colored according to the k-means cluster assignment.

**Supplementary Figure 3.**
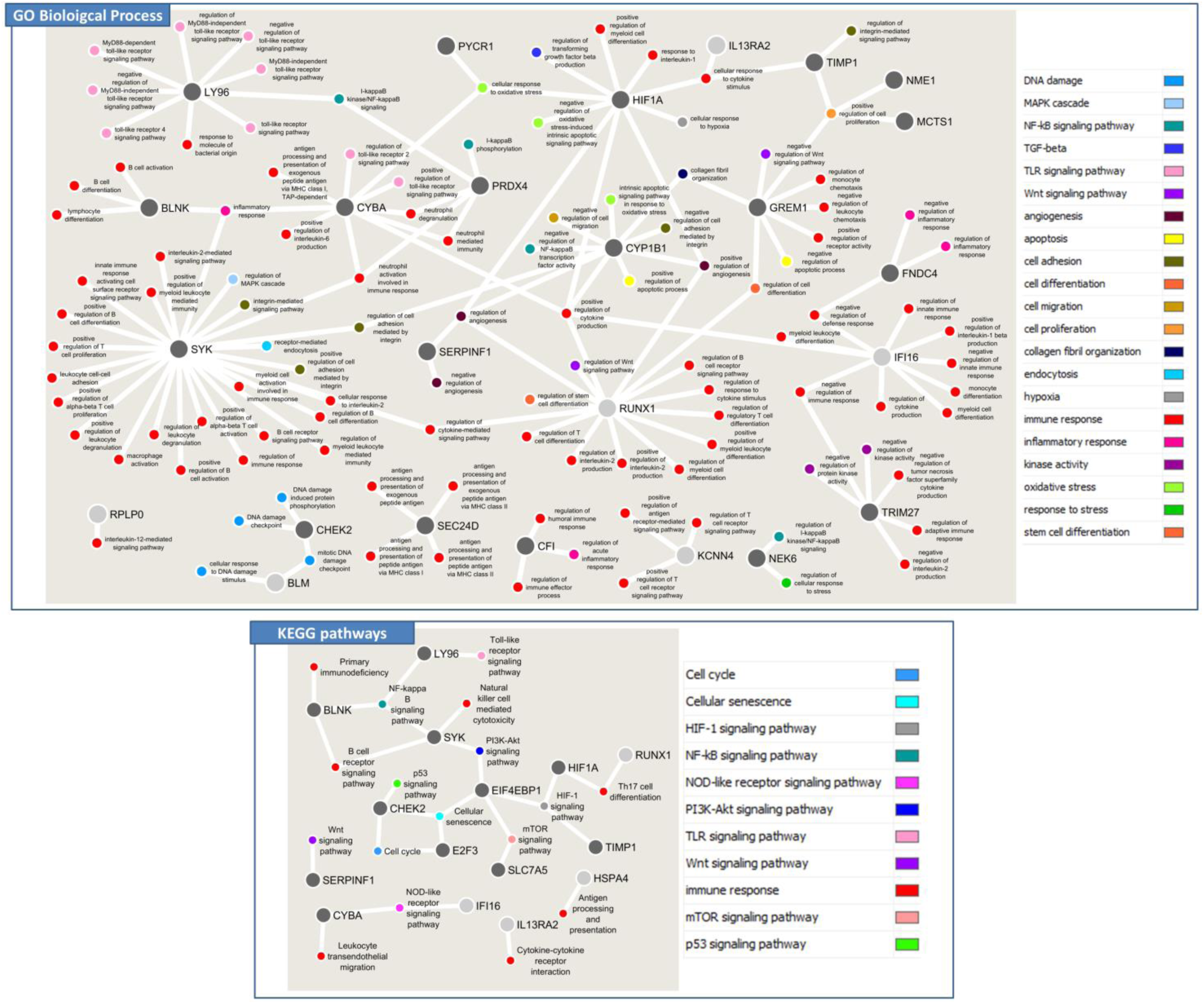
Functional annotation network of switch genes. In both panel larger dots represent switch genes, smaller dots represent the GO terms (top) and KEGG pathways (bottom) in which they are involved. GO terms and KEGG pathways are coloured according different categories reported in the legend. Switch genes nodes are coloured according to their cluster belonging (i.e., light grey = module 2; dark grey = module 3).

**Supplementary Figure 4.**
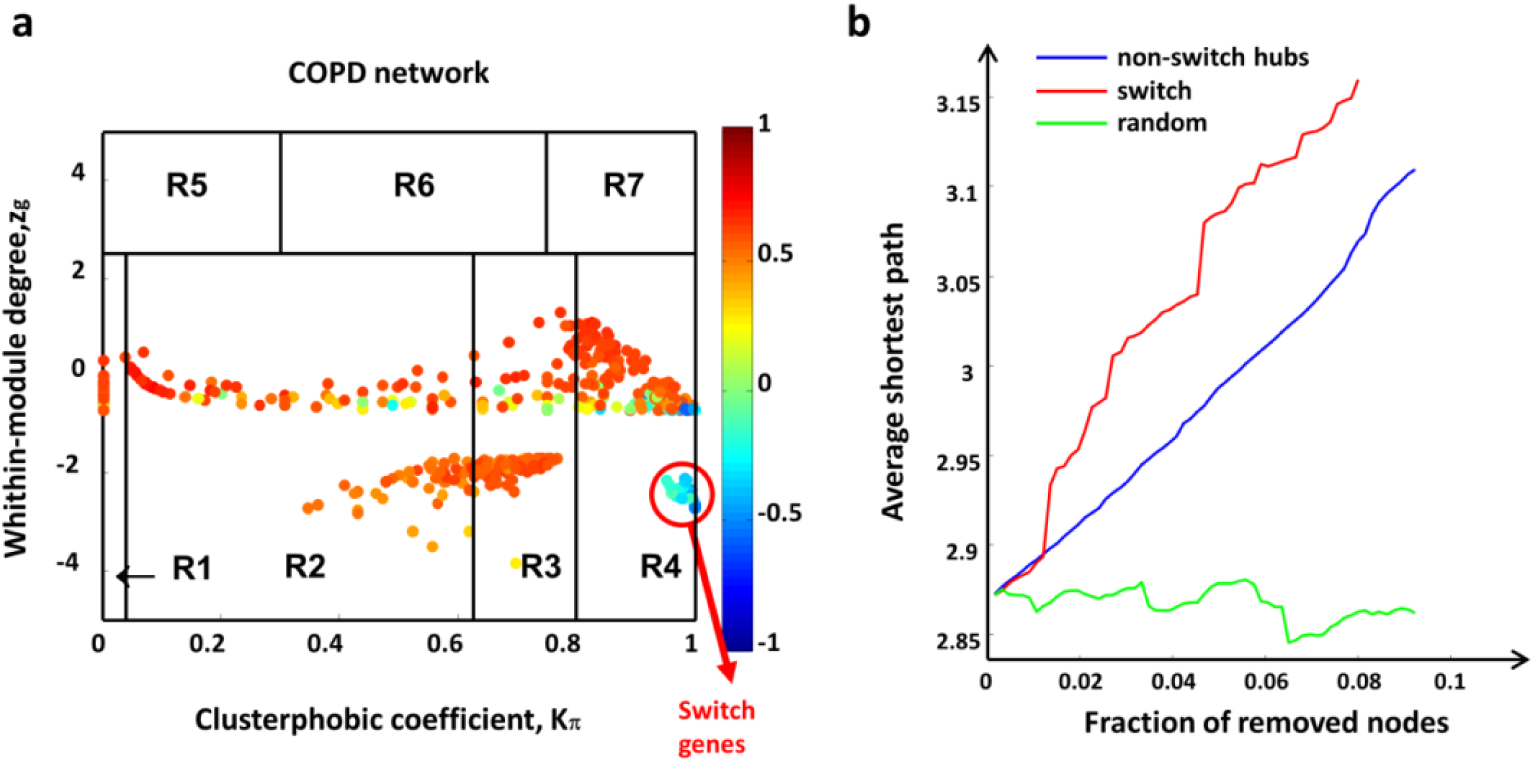
COPD switch genes for the test set (GSE76925 dataset). **(a)** Heat cartography maps of COPD network. Dots correspond to nodes in the networks. Each node is colored according to the value of the APCC between its expression profile and that of its nearest neighbors in the network. **(b)** Robustness for COPD correlation network. Blue curve corresponds to the cumulative deletion of non-switch hubs (i.e. the first 61 hubs that are not switch genes, sorted by decreasing degree); red curve corresponds to the cumulative deletion of the 61 switch genes, sorted by decreasing degree; the green curve corresponds to the cumulative deletion of randomly selected nodes. The x-axis represents the cumulative fraction of removed nodes with respect to the total number of network nodes that is 667 (i.e. x-maximum is 61/667 = 0.09), while the y-axis represents the average shortest path.

## Supplementary Tables

**Supplementary Table 1. COPD differentially expressed genes.** The table is composed of three separate sheets reporting: the statistically significant differentially expressed genes obtained for the COPD training set (GSE47460 dataset) along with their statistics (sheet 1); the differentially expressed genes of the COPD training set (GSE47460 dataset) that were previously identified as genome-wide significant COPD GWAS genes (sheet 2); and the statistically significant differentially expressed genes obtained for the COPD test set (GSE76925 dataset) along with their statistics (sheet 3).

**Supplementary Table 2. Node characterization in the COPD co-expression network.** The table is composed of four separate sheets reporting: all network nodes obtained for the COPD training set (GSE47460 dataset) along with their attributes and their statistics (sheet 1); bar plots reporting the number of nodes of COPD network for training set in each hub category and in each cluster (sheet 2); table reporting the immune cell-related genes for six different immune cell types (i.e., B cells, Macrophages, Monocytes, Neutrophils, Plasma cells, T cells) found in the network modules of COPD training set (sheet 3); all network nodes obtained for the COPD test set (GSE76925 dataset) along with their attributes and their statistics (sheet 4).

**Supplementary Table 3. Module membership.** The table is composed of three separate sheets reporting the list of genes belonging to each COPD network module for the training set (GSE47460 dataset) along with their module membership and their statistics.

**Supplementary Table 4. COPD switch genes.** This table is composed of six separate sheets reporting: the list of switch genes identified by SWIM for the COPD training set (GSE47460 dataset) along with their attributes and their statistics (sheet 1); the list of switch genes identified by SWIM for the COPD test set (GSE76925 dataset) along with their attributes and their statistics (sheet 2); KEGG pathways and GO Biological Process found to be associated with switch genes of the COPD training set (sheet 3-4); KEGG pathways and GO Biological Process shared between the switch genes of the COPD training set and the switch genes of the COPD test set (sheet 5-6).

**Supplementary Table 5. Nearest neighbors of COPD switch genes.** This table is composed of four separate sheets reporting for the COPD training set (GSE47460 dataset): the list of switch genes with their positively correlated GWAS genes along the columns for the COPD training set (sheet 1); the list of switch genes with their negatively correlated GWAS genes along the columns (sheet 2); the list of switch genes which negatively correlate with the GWAS genes AGER (sheet 3) and EMP2 (sheet 4).

